# Behavioral screening of conserved RNA-binding proteins reveals CEY-1/YBX RNA-binding protein dysfunction leads to impairments in memory and cognition

**DOI:** 10.1101/2024.01.05.574402

**Authors:** Ashley N Hayden, Katie L Brandel, Paul R Merlau, Priyadharshini Vijayakumar, Emily J Leptich, Edward W Pietryk, Elizabeth S Gaytan, Connie W Ni, Hsiao-Tuan Chao, Jill A Rosenfeld, Rachel N Arey

## Abstract

RNA-binding proteins (RBPs) regulate translation and plasticity which are required for memory. RBP dysfunction has been linked to a range of neurological disorders where cognitive impairments are a key symptom. However, of the 2,000 RBPs in the human genome, many are uncharacterized with regards to neurological phenotypes. To address this, we used the model organism *C. elegans* to assess the role of 20 conserved RBPs in memory. We identified eight previously uncharacterized memory regulators, three of which are in the *<u>C. e</u>legans* <u>Y</u>-Box (CEY) RBP family. Of these, we determined that *cey-1* is the closest ortholog to the mammalian <u>Y</u>-<u>B</u>o<u>x</u> (YBX) RBPs. We found that CEY-1 is both necessary in the nervous system for memory ability and sufficient to increase memory. Leveraging human datasets, we found both copy number variation losses and single nucleotide variants in *YBX1* and *YBX3* in individuals with neurological symptoms. We identified one predicted deleterious *YBX3* variant of unknown significance, p.Asn127Tyr, in two individuals with neurological symptoms. Introducing this variant into endogenous *cey-1* locus caused memory deficits in the worm. We further generated two humanized worm lines expressing human *YBX3* or *YBX1* at the *cey-1* locus to test evolutionary conservation of *YBXs* in memory and the potential functional significance of the p.Asn127Tyr variant. Both *YBX1/3* can functionally replace *cey-1*, and introduction of p.Asn127Tyr into the humanized *YBX3* locus caused memory deficits. Our study highlights the worm as a model to reveal memory regulators and identifies YBX dysfunction as a potential new source of rare neurological disease.

**GRAPHICAL ABSTRACT:** 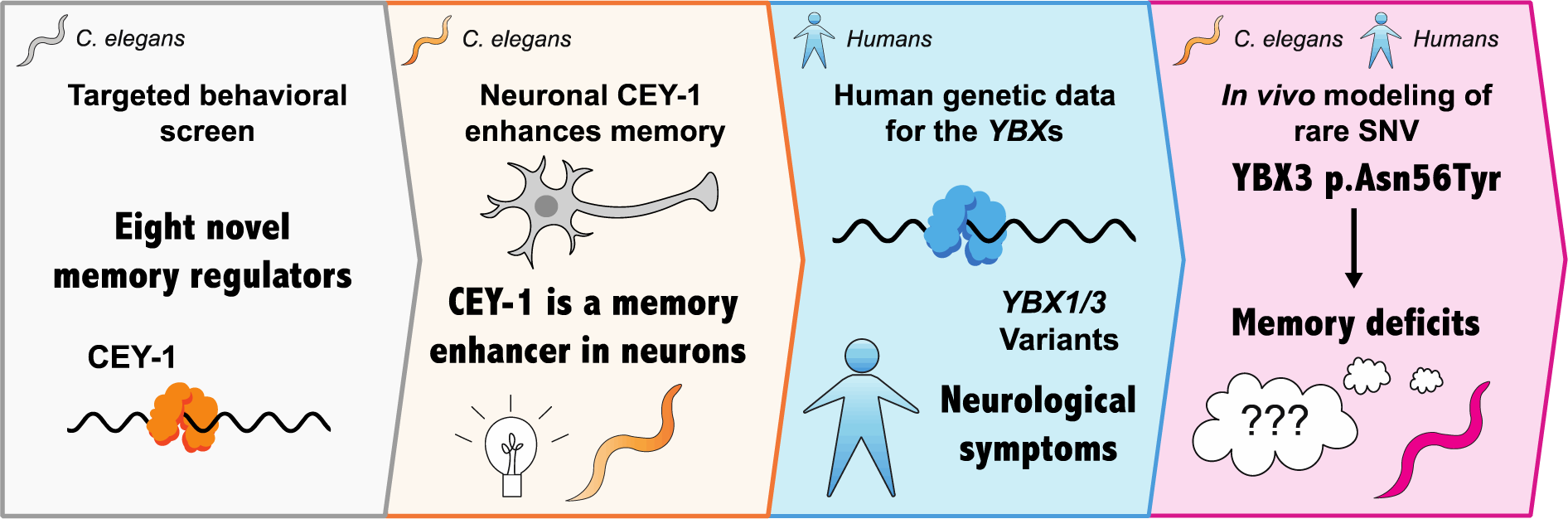

## INTRODUCTION

Careful regulation of RNA localization, translation, and stability is essential for the proper functioning of all tissues but is especially important in the nervous system. Neurons are composed of subcellular compartments, specifically somatic, axonal, and dendritic regions, each of which possesses a specialized and functionally distinct proteome. Moreover, these subcellular compartments must respond dynamically, and often uniquely, to external stimuli. A prominent example of this is local mRNA translation in response to plasticity-inducing stimuli, which is necessary for learning and memory [1,2]. In this instance, both the pre- and post-synaptic compartment can undergo local protein synthesis in response to the same plasticity-inducing stimulus [3,4], yet mRNAs translated in response to that stimulus are compartment-specific, likely due to distinct mRNA regulators.

RNA-binding proteins (RBPs) are involved in each step of RNA regulation in neurons, controlling target RNA splicing, polyadenylation, localization, translation, and stability [5]. As such, their proper function is essential for neuronal plasticity and its associated behaviors – especially learning and memory. The importance of understanding the role of RBPs in the nervous system is underscored by a growing body of evidence that their dysfunction results in neurological disease, including disorders ranging from neurodevelopmental disorders to neurodegenerative disease [6–8]. A frequent symptom of RBP-associated disorders are behavioral abnormalities and cognitive impairments, including Fragile X Messenger Ribonucleoprotein (FMRP) in Fragile X syndrome involving cognitive impairment, PUMILIO1 (PUM1) associated developmental disability, ataxia, and seizures (PADDAS), Quaking (QKI) in schizophrenia, and both Heterogeneous Ribonucleoprotein K (HNRNPK) in Au-Kline syndrome and SON in Zhu-Tokita-Takenouchi-Kim (ZTTK) syndrome both involving intellectual disability [9–13]. In fact, RBPs outweigh all other protein types in their prevalence of predicted deleterious variation in Mendelian disorders, especially those that are diseases of the nervous system [7]. However, despite their importance, many RBPs are unexamined in the nervous system. Barriers remain to systematic characterization of RBPs, as such studies remain relatively slow in mammals and the human genome encodes nearly 2000 RBPs, which would require extensive time and effort to comprehensively study [14,15]. Moreover, studying the tissue-specific function of RBPs is even more difficult as it requires challenging and costly genetic approaches. However, these issues can be circumvented in simpler model organisms like the nematode worm *Caenorhabditis elegans (C. elegans)* that allow for low-cost, rapid screening of molecules in a tissue-specific manner.

*C. elegans* have a rich history of use in high-throughput approaches and led to the discovery of many new regulators of neurological phenotypes, including molecularly conserved behaviors [16–18]. Indeed, the use of the worm resulted in the discovery of many conserved regulators of disease, including those involving cognitive deficits [19–24]. The *C. elegans* genome encodes almost 900 RBPs, over 80% of which have mammalian orthologs [25], and can be studied using thousands of publicly available mutants, including those that allow for neuron-specific RNAi for rapid tissue-specific genetic screening [26,27]. Importantly, *C. elegans* offers the advantage of using these genetic tools to investigate the role of RBPs in memory and cognition, as they form molecularly conserved associative memories, including short-term (STM), intermediate-term (ITM), and long-term memories (LTM) [20,28–32]. Given its wealth of molecular tools, ease of genetic screening, and molecularly conserved memory, *C. elegans* is an excellent model system to define the functions of neuronal RBPs in learning and memory *in vivo*.

Here, we have used the worm to perform a neuron-specific, targeted screen to discover conserved RBPs that control memory. Altogether, we find that 80% of our screened RBPs are memory regulators, eight of which are novel memory molecules. Importantly, our approach reveals a family of RBPs that are practically uncharacterized in their nervous system function and are essential for associative memory ability in the worm: the *<u>C</u>. <u>e</u>legans* <u>Y</u>-Box (CEY) RNA binding protein family.

Of this family, we determine that CEY-1 is most closely related to mammalian YBXs and requires further study to understand the potentially conserved role of YBXs in the nervous system. We found that CEY-1 acts specifically in the nervous system to promote memory. We discovered that this role in memory is likely conserved, as human copy number losses of *YBX* genes may be associated with neurological symptoms, most commonly intellectual disability. We similarly describe one rare heterozygous variant in the human *YBX3* gene identified in two unrelated individuals with neurological symptoms. Importantly, introduction of the conserved *YBX3* variant into either endogenous *cey-1* locus or animals where the *cey-1* locus has been humanized causes memory deficits, supporting the idea that deleterious *YBX3* variants may contribute to human disease.

Altogether, we uncovered new associative memory regulators in an unbiased screen of conserved neuronal RBPs. We describe in detail a novel associative memory regulator, CEY- 1/YBX, and highlight *C. elegans* as a discovery organism for associative memory regulators and potential disease-causing RBPs.

## Materials and Methods

### *C. elegans* maintenance

All strains were maintained at 20°C on 10cm plates made from standard nematode growth medium (NGM: 3g/L NaCl, 2.5g/L of Bacto-peptone, 17g/L Bacto-agar in milliQ water) or high nematode growth medium (HGM: 3g/L NaCl, 20g/L of Bacto-peptone, 30g/L Bacto-agar in milliQ water). After autoclaving and allowing molten agar to cool slightly, we added 1mL/L cholesterol (5mg/mL in ethanol), 1mL/L 1M CaCl_2_, 1mL/L 1M MgSO_4_, and 25mL/L 1M potassium phosphate buffer (pH 6.0) [33]. Experiments were performed using NGM plates seeded with OP50 *E. coli* as the food source for *ad libitum* feeding [33].

Hypochlorite population synchronization was performed by collecting eggs from gravid hermaphrodites via exposure to an alkaline-bleach solution (85mL water, 15mL sodium hypochlorite, 5mL 5M NaOH), followed by repeated washing of collected eggs in 1mL of M9 buffer (6g/L Na_2_HPO_4_, 3g/L KH_2_PO_4_, 5g/L NaCl and 1mL/L 1M MgSO_4_ in milliQ water [33]).

### Strains

Wild-type: (N2 Bristol)

Mutants: NM1380(*egl-30(js126)),* LC108(*uIs69[punc-119::sid-1; pmyo-2::mCherry]),* TU3401*(usIS69[myo-2p::mCherry + unc119p::sid-1]; sid-1(pk3321))*, RAF5(*cey-1(rrr12)),* VC1310*(cey-1(ok1805)),* GLW51*(cey-1(utx43[mNG::3xFLAG::cey-1]) II),* OH13605*(otIs619[unc- 11(prom8)::2xNLS::TagRFP] X)* were obtained from the Caenorhabditis Genetics Center (University of Minnesota, Minneapolis, MN).

Nuclear GFP reporter for *cey-1* #1087*(cey-1p::PEST::GFPH2B::cey-1u; unc-119 was* described previously [34] and generously gifted by Rafal Ciosk.

Strains made in collaboration with InVivo Biosystems include: COP2655*(knuSi967[(rab-3p::CEY- 1::3xFLAG::rab-3u, unc-119(+))] IV; unc-119(ed3) III)* and COP2656*(knuSi968[(rab-3p::CEY- 1::3xFLAG::rab-3u, unc-119(+))] IV; unc-119(ed3) III)* are two lines with the same genotype, COP2689(*cey-1(knu1233 [N56Y]));* COP2721*((knu1251[YBX3]) II);* COP2763((*knu1275[YBX1]) II*); COP2774((*knu1286[p.N127Y]) II*) and COP2775((*knu1287[p.N127Y]*) II) are two lines with the same genotype.

The following strains were generated by crosses: RNA1*(egl-30(js126); uIs69[(pCFJ90)myo- 2p::mCherry + unc-119p::sid-1])* was made by crossing NM1380*(egl-30(js126))* with LC108*(uIs69[(pCFJ90)myo-2p::mCherry + unc-119p::sid-1]).* RNA20(*cey- 1(utx43[mNG::3xFLAG::cey-1]) II; otIs619[unc-11(prom8)::2xNLS::TagRFP] X)* was made by crossing OH13605*(otIs619[unc-11(prom8)::2xNLS::TagRFP] X)* with GLW51*(cey- 1(utx43[mNG::3xFLAG::cey-1]) II).* RNA26(*cey-1p::PEST::GFPH2B::cey-1u; unc-119(+); otIs619[unc-11(prom8)::2xNLS::TagRFP] X)* was made by crossing OH13605*(otIs619[unc- 11(prom8)::2xNLS::TagRFP] X)* with *cey-1* #1087*(cey-1p::PEST::GFPH2B::cey-1u; unc-119(+)).* RNA40*(cey-1(rrr12); knuSi968[(rab-3p::CEY-1::3xFLAG::rab-3u, unc-119(+)] IV; unc-119(ed3) III)* was made by crossing RAF5(*cey-1(rrr12))* with COP2656*(knuSi968[(rab-3p::CEY-1::3xFLAG::rab- 3u, unc-119(+))] IV; unc-119(ed3) III)*.

### RNAi treatment

For adult-only RNAi, worms that are neuronally sensitive to RNAi, RNA1*(egl-30(js126); uIs69[(pCFJ90)myo-2p::mCherry + unc-119p::sid-1])),* or allow RNAi only in the nervous system, TU3401*(usIS69[myo-2p::mCherry + unc119p::sid-1]; sid-1(pk3321)),* were synchronized by bleaching. Then, at L4, worms were transferred from plates seeded with OP50 to plates seeded with RNAi (*HT115 E. coli)* for *ad libitum* feeding and allowed to grow for two days. For all RNAi experiments, standard NGM molten agar was supplemented with 1mL/L IPTG (isopropyl ꞵ-d-1-thiogalactopyranoside) and 1mL/L 100mg/mL carbenicillin.

### Short-term and intermediate-term associative memory assays

Wild-type, mutant, or RNAi-treated worms were trained and tested for short- and intermediate-term memory changes as previously described [28]. Briefly, synchronized Day 2 adult worms were washed off plates with M9 buffer. Worms were then allowed to settle by gravity and washed twice more with M9 buffer to remove any bacteria. After washing, the worms were starved for one hour in M9 buffer. For 1x food-butanone pairing, hereby called conditioning, starved worms were transferred to 10cm NGM conditioning plates seeded with OP50 *E. coli* bacteria and with a total of 16μL of 10% butanone (Sigma Aldrich) diluted in ethanol streaked on the lid in a ‘#’ shape for one hour. After conditioning, the trained population of worms were tested for chemotaxis to 10% butanone and to an ethanol control using standard, previously described chemotaxis conditions [35]. Different stages of memory were tested by measuring chemotaxis of different subpopulations of worms at different timepoints at molecularly distinct stages of memory [29,30]. These stages are immediately after training (0min, learning) or after being transferred to 10cm NGM plates with fresh OP50 for 30 minutes (short-term associative memory), 1 hour (intermediate term associative memory), or 2 hours (forgetting).

Chemotaxis indices for each timepoint were calculated as

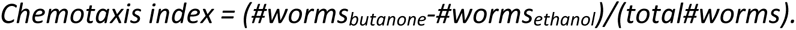

Performance index is the change in the chemotaxis index after training relative to the untrained chemotaxis index, or

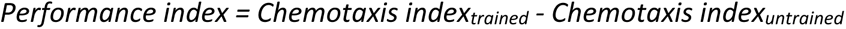

### Long-term associative memory assays

As previously published, *egl-30(js126)* animals form a long-term memory after just one round of training [36]. For LTM screening, we administered RNAi to adult, neuronally RNAi sensitive *egl- 30(js126);punc119::sid-1* worms, performed S/ITM training (1 CS-US pairing), and measured learning immediately after training and long term memory 16-20 hours post-training. Chemotaxis index and performance index were calculated in the same manner as short term and intermediate term associative memory assays.

### Baseline chemosensation assays

Chemotaxis, or chemosensation experiments, were performed based on previously published assays [35]. In brief, assays were performed on unseeded 10cm NGMs. Two marks were made on the back of the plate on opposite sides of the plate, approximately 0.5cm from the edge. 1μL of sodium azide (Thermo Fisher) was placed on both spots and allowed to dry before adding 1μL of test odorant diluted in ethanol on one side and ethanol on the other. Odorants included 0.1% and 10% butanone (vol/vol), 0.1% nonanol (2-nonanone)(vol/vol), 1% isoamyl alcohol (vol/vol), 1% benzaldehyde (vol/vol), 10% pyrazine (weight/vol), and 1% diacetyl (vol/vol) (all from Sigma Aldrich). Worms were washed off their plates and subsequently washed three times with M9 buffer, then placed near the bottom center of the plate, equidistant between the two marks, and allowed to chemotax for an hour. Chemotaxis indices for each timepoint were calculated as

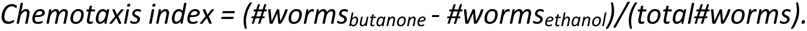

### Motility assays

We measured motility as previously published [20] by finding the average percentage of worms remaining at the origin of the plate after a naïve (untrained) chemotaxis assay using an attractive concentration of butanone (0.1% vol/vol).

### Confocal Microscopy

Worms were paralyzed with fresh 4% levamisole diluted in M9 buffer and imaged for up to 30 minutes once put onto the slide to preserve protein dynamics. In all experiments, imaging of Day 2 adult worms was performed on a Nikon Ti2E inverted microscope system with a W1 spinning disk confocal unit at 100x magnification. We used an excitation wavelength of 488nm for GFP and 561nm for RFP, used a pixel size of 0.11um/px, and z stacks with a z-step of 0.6nm. Images were processed in Nikon NIS Elements software. The same settings for laser power and detector gain were used for all genotypes. For all images, red fluorescence was pseudo colored magenta and green fluorescence were pseudo colored cyan for colorblindness inclusivity.

### Variant identification and pathogenicity analysis

We received a deidentified dataset of human *YBX* gene variants identified through clinical exome and genome sequencing completed at Baylor Genetics [37] and removed any variants where symptomology listed could be explained by other genetic alterations found in the individual’s genome. Next, we removed any variants found in gnomAD v.2.1.1, even if only one case was reported [38], as these variants would not likely cause severe disease if found in the general population. We then selected variants for potential *in vivo* modeling based on the presence of neurological symptoms, such as seizures, microcephaly, autism, and intellectual disability. Variants of interest were then assessed using multiple pathogenicity metrics to determine the final variant for *in vivo* testing: specifically, M-CAP [39], SIFT [40], PolyPhen 2 [41], GERP [42], CADD [43,44], REVEL [45], and Phylop Vertebrate [46]. All metrics were found and visualized using the UCSC genome browser [47]. We selected the variant with the highest overall predicted pathogenicity across all score metrics as opposed to any specific numerical cutoffs.

### Phylogenetic analysis

To analyze the evolutionary history and determine the similarity of RNA binding proteins between phyla, we used maximum likelihood (ML) analysis with the JTT matrix-based model [48]. We obtained protein sequences of the *C. elegans* CEY proteins as well as the human YBX proteins from UniProt [49] then used these sequences to generate a tree with the highest log likelihood score (-4517.57) using the Neighbor-Join and BioNJ algorithms to calculate pairwise distances from the JTT model. The tree was drawn to scale, with branch lengths measured in the number of substitutions per site and a 0.20 scale bar indicating the relative distance between nodes. All analyses were performed using MEGA X [50,51].

### Orthology analysis

Drosophila RNAi Screening Center Integration Ortholog Prediction Tool (DIOPT) is an online bioinformatics resource designed to facilitate the identification of orthologous genes across various species [52].

### RNA isolation, cDNA synthesis and qRT-PCR

Worms of a particular genotype were crushed in liquid nitrogen and added to Trizol (Thermo Fisher Scientific). RNA was isolated per manufacturer’s instructions, followed by DNase treatment (Qiagen). cDNA was synthesized with an oligo dT primer and Superscript III reverse transcriptase enzyme (Thermo Fisher Scientific). cDNA was mixed with buffers, primers, SYBR green, and hot start Taq polymerase in a master mix prepared by a manufacturer (Thermo Fisher Scientific). Using a Quant Studio 7 Pro Dx Real-Time PCR System (Thermo Fisher Scientific), PCR reactions were run followed by a dissociation reaction to determine specificity of the amplified product. The amount of gene expression was quantified using the ΔΔCt method using pmp-3 as a reference gene. Primer sets were as follows: *cey-1* For: 5’-GGATCCAAGTATGCTGCCGA -3’ *cey-1* Rev: 5’- CCATCTGTGTCACGAGCAGT -3’ *pmp-3* For: 5’- AGTTCCGGTTGGATTGGTCC -3’ *pmp-3* Rev: 5’- CCAGCACGATAGAAGGCGAT-3’

### Statistical analysis

Statistical data is reported in the main text, figures, and tables as noted. Significance threshold of p < 0.05 was used. The symbols *, **, ***, and **** refer to p < 0.05, 0.01, 0.001, and 0.0001, respectively. For the comparison of performance indices between two behavior conditions (eg. vector control vs *cey-1* RNAi), a Mann-Whitney test comparing ranks was used because it does not assume normality. For comparison of performance indices between three or more groups (eg wild-type vs two different *cey-1* loss-of-function mutants), one-way analysis of variances followed by Bonferroni post hoc tests for multiple comparisons were performed.

Two-way ANOVAs were used for evaluating effects between genotype (Wild-type, *cey-1(rrr12)*, *YBX1*, *YBX3*) and timepoint (0hr, 0.5hr, 1hr, 2hr) on performance indices with a significant interaction between factors (p<0.0001) prompting Bonferroni post-hoc analyses to determine differences between individual groups. All experiments were repeated on separate days with separate populations to confirm reproducibility of results. Sample size n represents the number of chemotaxis assays performed for behavior, with each assay containing approximately 50-150 worms each.

### Statistical analysis software

All statistics and code were run in GraphPad Prism 10, using standard toolboxes.

## RESULTS

### Targeted screen identifies novel RBPs that regulate learning and memory

The *C. elegans* genome, which exhibits a high degree of conservation with mammals, encodes 887 predicted RBPs [53,54]; thus, the worm is an excellent system to rapidly screen RBPs for conserved roles in memory. We prioritized candidate RBPs based on the following criteria to increase the likelihood of identifying memory regulators: 1) nervous system expression based upon transcriptomic studies, 2) evolutionary conservation with mammals, and 3) detection in synaptic regions across species, as RBPs are known to regulate plasticity at the synapse. By cross-referencing adult only, neuron specific transcriptomic datasets [24,55] as well as orthology tools including Wormcat and OrthoList2 [56,57], we identified 550 neuronally expressed RBPs with mammalian orthologs. After filtering this list based upon detection in *C. elegans* and mammalian synapses [55,58–62], 313 RBPs met screening criteria. Of those, we proceeded with the top 20 most synaptically-enriched RBPs based on our previous work identifying the adult *C. elegans* synaptic transcriptome [58,63] for behavioral study.

We assessed the roles of the RBPs in memory by performing adult-only RNAi in neuronally RNAi-sensitized animals *(punc119::sid-1)* [27] and measuring the effects of knockdown on various well-characterized, molecularly conserved associative behaviors. These behaviors included learning (0 min after training), protein synthesis-independent short term memory (STM, 30 min after training), and translation-dependent intermediate term memory (ITM, 1 hr after training) (**Figure 1A**) [29,30]. Effects of RBP knockdown on transcription and translation-dependent long term memory (LTM) were assessed in RNAi-sensitized animals with an additional *egl-30* gain of function mutation *(egl-30(js126), punc119::sid-1))* that increases G_αq_ signaling and confers the ability to form a LTM that is identical to wild-type animals with only one round of training, making them amenable to high-throughput screening (**Figure 1B**) [36].

**Figure 1:**
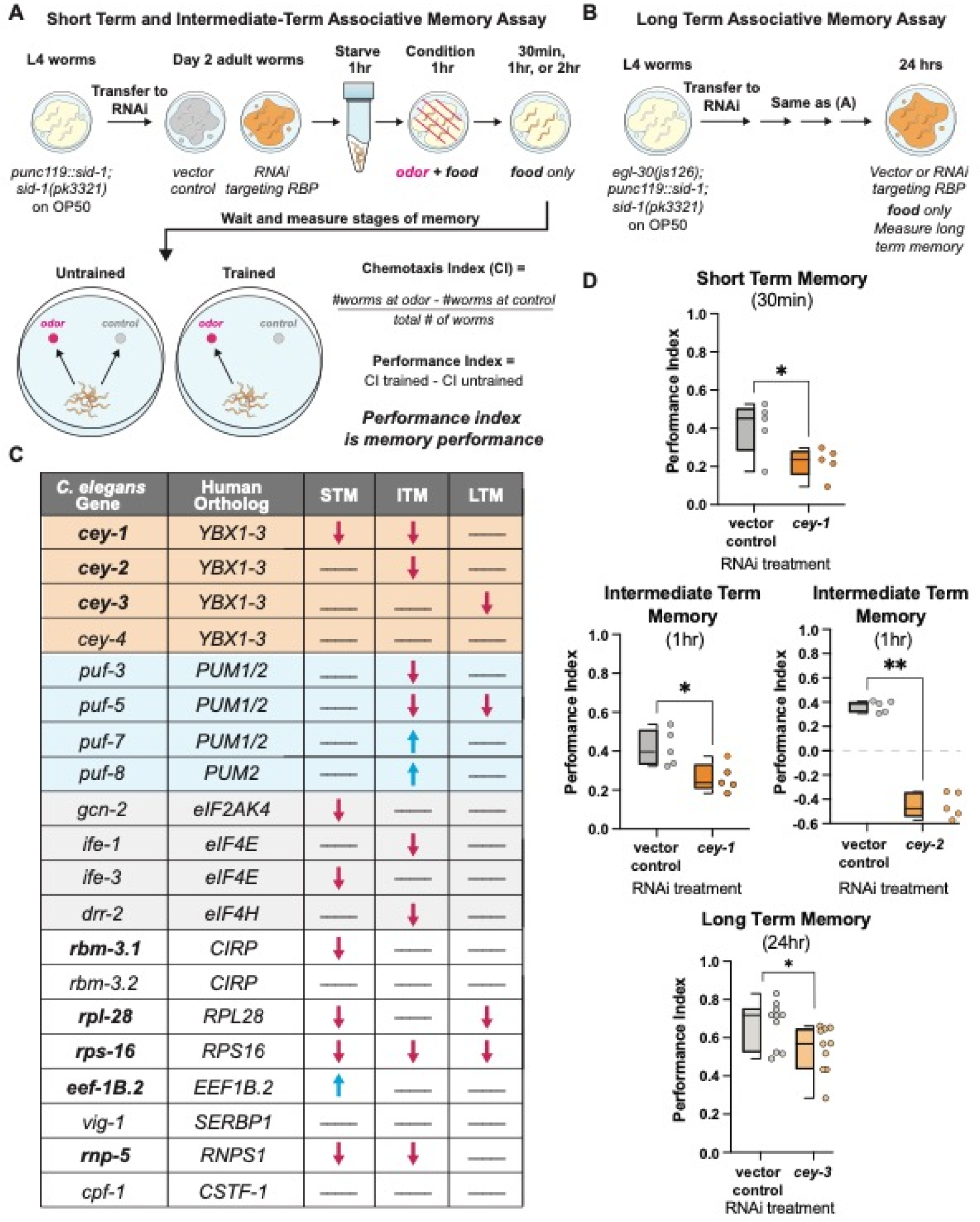
Unbiased screen reveals novel associative memory regulators. (**A**) Short term/intermediate term associative memory (STM/ITM) assay workflow using adult-only RNAi knockdown [29,30]. Adult worms are starved for 1 hour, conditioned for 1 hour, and tested immediately for 1x learning (0 min) via chemotaxis assays, or transferred onto holding plates with OP50 for 0.5, 1, or 2 hours after conditioning, then tested for STM and ITM changes. Memory performance is calculated by comparing the trained and untrained chemotaxis indices. (**B**) The long-term associative memory (LTM) assay workflow is identical to that for STM/ITM except worms have an additional *egl-30(js126)* mutation that allows for LTM formation after just 1x training [36]. Worms are tested immediately for 1x learning (0 min) via chemotaxis assays or transferred onto holding plates with RNAi for 16-24 hours after conditioning, then tested for LTM changes. (**C**) Results of unbiased screen of 20 RBPs reveals eight novel associative memory regulators. All RBPs screened are shown and include the *C. elegans* gene, its mammalian ortholog, and any STM, ITM, or LTM changes. Blue arrows indicate enhanced memory while red arrows indicate decreased memory; lines indicate memory did not change. Novel memory regulators not reported in the literature are bolded. For full screen results, see **Figure S1-2.** (**D**) Each member of the CEY RBP family plays a novel, unique role in memory. Adult-only RNAi knockdown of *cey-1* in neuronally-sensitized worms causes STM and ITM deficits, *cey-2* knockdown causes ITM deficits, and *cey-3* knockdown causes LTM deficits. Box and whisker plot: the center line denotes the median value (50th percentile) while the box contains the 25th to 75th percentiles. Whiskers mark the 5th and 95th percentiles.

Of the 20 RBPs tested, 80% control at least one type of memory (**Figure 1C**). Two classes of proteins known to control memory across species had multiple members represented in our screen candidates: the Pumilio and FBF RBP family (*pumilio1/2* orthologs *puf-3*, *puf-5*, *puf-7*, and *puf-8*) as well as four proteins involved in translation initiation (eIF4/2α orthologs *gcn-2, ife-1, ife-3,* and *drr-2*). Pumilios have been linked to both LTM and working memory in *Drosophila melanogaster* and mice [64,65], and we have previously described their importance in *C. elegans* memory [58,63]. Similarly, eukaryotic translation initiation machinery eIF4E and eIF2α control memory formation, STM, and LTM in mice [66,67]. However, to the best of our knowledge the requirement for this molecular machinery in olfactory associative memory was not previously demonstrated in the worm. Validating these two classes of known memory regulators strengthens the likelihood that novel molecules identified in our screen have a conserved function in higher organisms. Indeed, we found that 50% of memory regulating RBPs in our screen were not previously associated with memory phenotypes, and are therefore novel, bolded in **Figure 1C**. Importantly, no learning deficits were detected after knockdown of any RBP in our screen (**Figure S1**), suggesting the deficits we identified are specific to memory and are not due to broad behavioral or neurological disruption. For all behavior timepoints where RNAi significantly altered memory, see **Figure S2.**

### Identification of CEY family as novel memory regulators

Strikingly, all members of the *<u>C. e</u>legans* <u>Y</u>-Box (CEY) RBP family (*cey-1-4*), orthologs of the mammalian *Y-box binding* (*YBX*) family, met our screening criteria despite being uncharacterized in learning and memory behavior. The CEY RBPs regulate fertility in the worm [34,68], and mammalian YBX proteins are translational regulators involved in cancer metastasis and cell cycle progression [69–72]. Here we found that three of the four CEY proteins screened regulate memory, with each CEY controlling its own molecularly distinct form of memory (**Figure 1D**); *cey-1* is required for STM and ITM, *cey-2* is required for ITM, *cey-3* is required for LTM, and *cey-4* knockdown had no detectable effect on behavior. To our knowledge, this is the first documented behavioral phenotype regulated by CEY RBPs.

As there are no reported neuronal phenotypes associated with the CEY RBPs, we further examined their expression pattern based on previous transcriptomic profiling datasets. Single-cell RNA-seq data generated from neurons isolated from L4 larval animals revealed broad expression of *cey-1* and *cey-4,* while *cey-2* is only detected in eight neurons (**Figure S3**)[73,74]. *Cey-3* is not detected in any neurons at L4 but is detected in adult neuron-specific datasets [24,60], highlighting the importance of compiling multiple transcriptomic datasets to choose our screen candidates.

Similar to their *C. elegans* orthologs, the mammalian YBX proteins are relatively understudied with regards to behavioral phenotypes or their function in the adult nervous system, though YBX1 is reported to be involved in neural stem cell development [75]. Similar to *cey-1*, *YBX1* is broadly expressed throughout the nervous systems of humans, adult macaques, and rats. In these mammals, *YBX1* is expressed in memory-regulating regions including the hippocampus, parahippocampal cortex, and dentate gyrus [76]. *YBX3* is also broadly expressed, including in memory-regulating regions, while *YBX2* is thought to be restricted to non-neuronal tissues [77,78]. Importantly, YBX1 is reported to bind to plasticity-associated mRNAs, including *GluR2* mRNA and *CaM1* mRNA, in an activity-dependent manner, making the YBXs promising candidates as novel essential memory molecules in mammals [79]. Moreover, both YBX1 and YBX3 proteins are detected in mouse hippocampal post-synaptic densities, suggesting that they have the potential to regulate local translation in a brain region critical for memory [80]. In non-neuronal tissues, CEYs and YBX proteins appear functionally conserved; both worm and mammalian proteins regulate polysome formation [34,81,82], suggesting that we can study conserved functions of this class of RBP in the worm.

To strengthen the translatability of our findings by focusing on conserved functions, we decided to study the *C. elegans* ortholog(s) that are the most closely related to the YBX proteins [71,72,75,83,84]. To this end, we performed phylogeny and homology analysis using MegaX and DIOPT [50,52] and found that *cey-1* is the primary *C. elegans YBX* ortholog, closer in similarity to the *YBXs* than other CEY family members (**Figure 2A, 2B**). For example, the linker regions (NC9 and CC13) and Cold-Shock Domain (CSD) of CEY-1 are extremely similar to those of the YBX proteins (approximately 97% similar) (**Figure 2C**) [85,86]. At both the N- and C-termini of the CEY-1/YBX proteins are moderately conserved (approximately 60%) low complexity regions, otherwise known as intrinsically disordered regions, responsible for protein-protein binding and stress granule formation [82,87].

**Figure 2:**
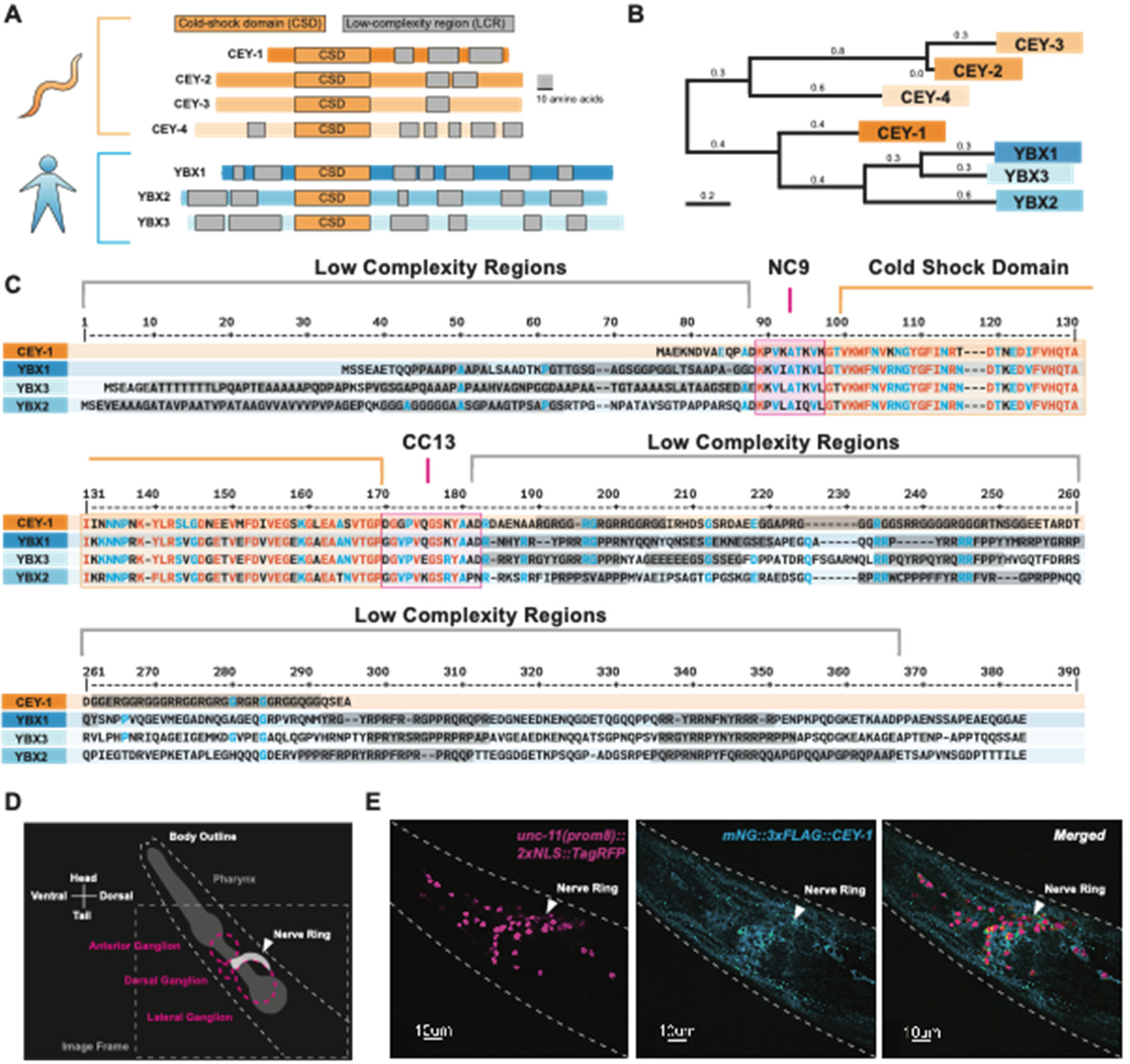
CEY-1 is the closest ortholog to the human Y-box binding proteins. (**A**) Diagram of the conserved protein domains of the *C. elegans* CEY RBPs and human YBX RBPs. *C. elegans* have four CEY RNA-binding proteins, CEY-1, CEY-2, CEY-3, and CEY-4, that are orthologs of the three mammalian Y-box binding proteins, YBX1, YBX2, and YBX3. All CEY/YBX RNA-binding proteins have a conserved cold-shock domain (CSD) that mediates RNA binding as well as one or more low complexity regions (LCRs) thought to be important for protein-protein interactions. Grey scale bar is 10 amino acids. (**B**) Phylogenetic tree generated using MegaX [50,51] reveals that CEY-1 is more similar to the human YBX proteins than the other CEY RNA-binding proteins. (**C**) Protein sequence alignment of the CEY and human YBX binding proteins made with MultiAlin [91] showing organization of protein domains. Highly conserved residues are depicted in red and blue. The cold-shock domains are highlighted in orange, the NC9 and CC13 linkers are highlighted in pink, and the low complexity regions are highlighted in grey. Domains were annotated according to SMART [92]. (**D**) Diagram of microscopy images shown in (E). *C. elegans* head is labeled including the location of the pharynx and main neuronal ganglia/nerve rings. (**E**) CEY-1 is primarily localized to the cytoplasm at baseline conditions. Representative image of Day 2 adult worms with RFP-labeled neuronal nuclei *(unc- 11(prom8)::2xNLS::TagRFP*) pseudocolored magenta and an endogenously mNeonGreen-tagged CEY-1 *(mNG::CEY- 1)* pseudocolored cyan.

Based on gene expression data, the expression pattern of *cey-1* and the mammalian YBX proteins is similarly broad across the nervous system [73,76–78,88–90]. Though previous studies using promoter-fusion approaches corroborate the transcriptomic evidence that *cey-1* is broadly expressed in the nervous system [34], we sought to verify that the protein is indeed expressed in neurons. We crossed the previously published *cey-1::GFP* promoter-reporter system with a marker of neuronal nuclei (*unc-11(prom8)::2xNLS::TagRFP*) and validated their reported neuronal expression (**Figure S4**). We confirmed these findings by verifying the neuronal localization of endogenous CEY-1 by crossing animals expressing an endogenously *mNeonGreen-tagged CEY-1 (mNG::CEY-1)* with *unc-11(prom8)::2xNLS::TagRFP* expressing animals (**Figure 2D,E**).

### CEY-1/YBX acts in neurons to promote memory

Despite its broad expression in the nervous system, we wanted to determine if *cey-1* was expressed in neurons involved in learning and memory. Because the *C. elegans* nervous system is invariant [93,94], the identities of specific neurons involved in a variety of associative behaviors are known. Based upon previously generated L4 single-cell RNA-seq data, we find that *cey-1* is indeed detected in all canonical memory-associated neurons, including those associated with gustatory plasticity, thermotaxis, and olfactory memory (**Figure 3A**) [35,36,95–100]. To confirm the finding from our initial screen that reduction of *cey-1* causes memory deficits, we examined the behavior of animals bearing two different loss-of-function *cey-1* alleles. Both animals exhibited severe deficits in learning and, as a result, inability to form a memory (**Figure 3B**).

**Figure 3:**
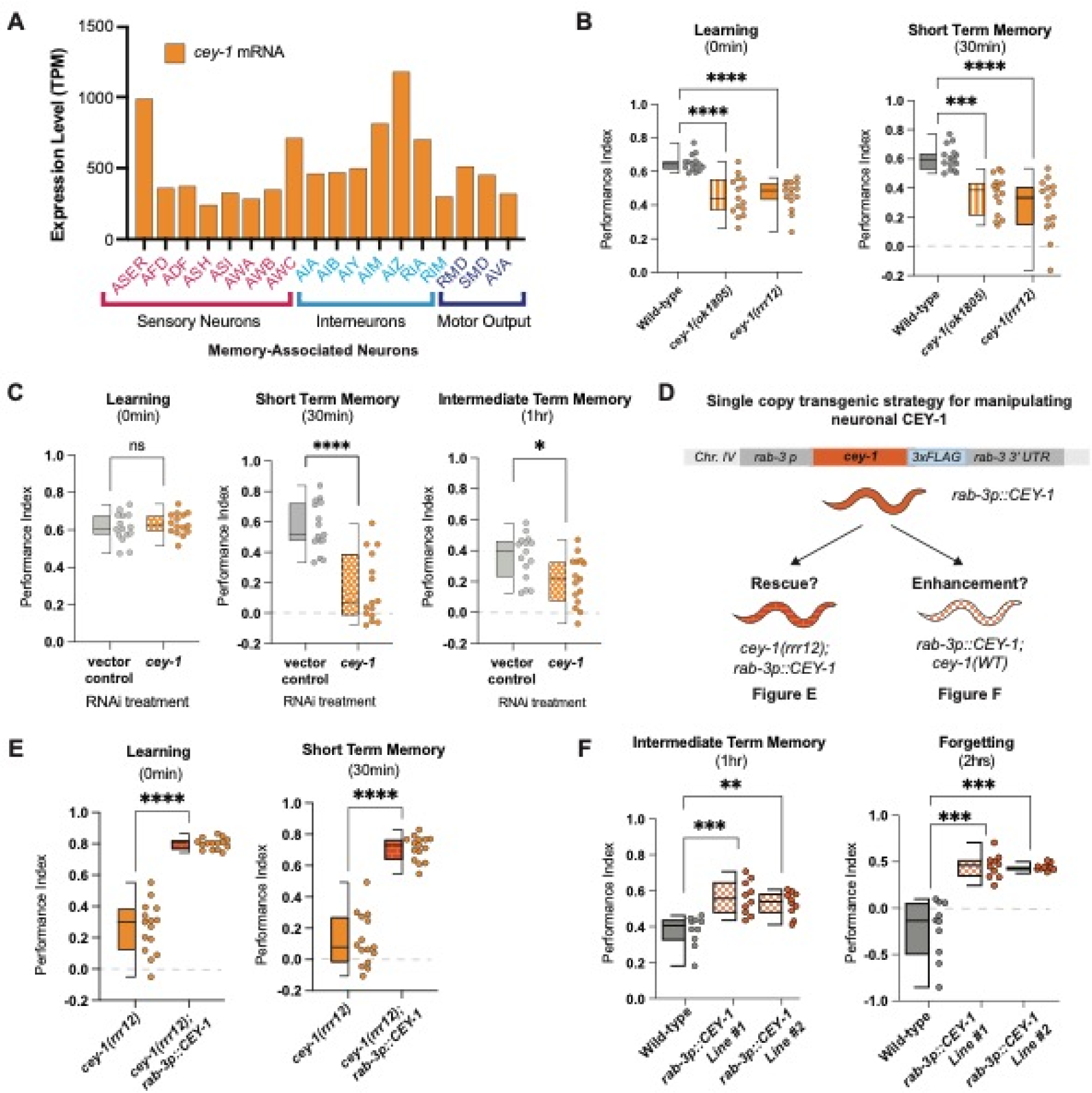
CEY-1 promotes memory in the *C. elegans* nervous system. (**A**) Single-cell RNA-seq data from L4 hermaphrodites[73] show that *cey-1* is expressed in every previously described memory-associated neuron [35,36,95–100]. (**B**) Loss-of-function mutants *cey-1(rrr12)* and *cey-1(ok1805)* have severe learning and memory deficits. Box and whisker plot as in Figure 1. n = 15 per genotype. ***p<0.001, ****p<0.0001. (**C**) Adult-only, neuron-specific knockdown of *cey-1* causes STM and ITM deficits. Box and whisker plot as in Figure 1. n = 15 per RNAi treatment. *p<0.05, ****p<0.0001. ns, not significant (p>0.05). (**D**) Diagram of strategy to determine whether neuronal *cey-1* is sufficient to rescue and enhance memory. Worms that express a single copy of *cey-1* only in the nervous system *(rab-3p::cey-1::rab-3 3’UTR; cey-1(rrr12))* were tested for phenotypic rescue in (E) while worms that have a single extra copy of *cey-1* only in the nervous system *(rab-3p::cey-1::rab-3 3’UTR; cey-1(+))* were tested in (F). (**E**) Single copy expression of *cey-1* in the nervous system *(rab-3p::cey-1::rab-3 3’UTR; cey-1(rrr12))* rescues loss-of-function learning and memory loss. Box and whisker plot as in Figure 1. n = 15 per genotype. ****p<0.0001. (**F**) Single copy overexpression of *cey-1* in the nervous system is sufficient to enhance memory *(rab- 3p::cey-1::rab-3 3’UTR; cey-1(+)).* Box and whisker plot as in Figure 1. n = 15 per genotype. **p<0.01,***p<0.001.

To determine if *cey-1* acts in the nervous system to control learning and memory, we next tested if nervous-system specific *cey-1* knockdown causes memory deficits by performing adult-only, neuron-specific knockdown of *cey-1* using worms only expressing *sid-1* in the nervous system *(punc119::sid-1; sid-1(pk3321)).* Contrary to the mutants used in our initial screen, which can have off-target effects in other tissues, these worms lack *sid-1* in all tissues other than the nervous system, leading to RNAi specifically in the nervous system [27]. Our findings replicated the initial results of our screen: *cey-1* knockdown decreases both STM and ITM but not learning (**Figure 3C**). Importantly, we detected no motility or butanone chemotaxis deficits (**Figure S5**), suggesting the deficits we observed are memory-specific. We believe that by knocking down *cey-1* only in neurons and during adulthood, we are circumventing loss of *cey-1* during development, which may be causing the more severe effects on learning seen in *cey-1* mutants.

### Elevated neuronal CEY-1 is sufficient to enhance memory

We next asked if restoring *cey-1* function in nervous system was sufficient to rescue learning and memory deficits observed upon loss of *cey-1*. We generated *cey-1(rrr12)* loss of function animals that also express a single-copy knock-in transgene driving expression of *cey-1* specifically in the nervous system *(rab-3p::cey-1::rab-3 3’UTR; cey-1(rrr12))* (**Figure 3D**). We found that neuron-specific *cey-1* rescue is sufficient to restore the learning and memory deficits seen in *cey-1(rrr12)* mutants back to wild-type levels (**Figure 3E, Figure S6**). Importantly, neither the *cey-1* loss of function mutants nor the neuronal *cey-1* rescue worms have butanone sensing deficits or impaired motility, suggesting that phenotypes recorded are memory-specific (**Figure S6**).

Thus far, our data suggested that *cey-1* acts in the nervous system to promote memory. To test this hypothesis, we determined whether increasing *cey-1* expression in the nervous system would have beneficial effects on behavior. A single round of food-butanone training results in memory that persists for no longer than two hours in wild-type animals, and we asked if additional neuronal *cey-1* increased the duration of this memory, as has been observed with other memory promoting genetic manipulations [28,29,63]. Worms with (*rab-3p::cey-1::rab-3 3’UTR; cey-1(+))* showed enhanced memory (**Figure 3F**), as their memory persists longer than the average two hours, consistent with previous memory enhancers [28,29,63]. Collectively, our results show that *cey-1* is sufficient to increase memory when expressed only in the nervous system, suggesting that *cey-1* is a memory enhancer.

### Copy number variant deletions in *YBX1* and *YBX3* may be associated with neurological symptoms, including intellectual disability

Human genomic datasets are increasingly linking RBP dysfunction to neurodevelopmental and neurological disorders, including those characterized by intellectual disability and cognitive dysfunction [6–8,101]. Therefore, we used copy number variation (CNV) datasets to determine if loss of the *YBXs* may potentially be associated with neurological dysfunction in humans, which would mirror our findings in *C. elegans*. We used an open-access database, DECIPHER, that reports CNVs from 50,000 individuals [102]. We focused specifically on CNVs in *YBX1* and *YBX3*, as *YBX2* is not expressed in the adult human brain [76,78].

We found that many individuals with CNV deletions that include either *YBX1* or *YBX3* have neurological symptoms (**Figure 4A**). The CNV deletions were variable in size, ranging from 1.59 Mb to 24.59 Mb. The most common symptom reported in individuals with CNV deletions including the *YBX*s is intellectual disability, as 37.5% of individuals with CNV deletions containing *YBX1* and 80% of individuals with CNV deletions in *YBX3* have intellectual disability (**Figure 4B**). Other common symptoms for individuals with *YBX1* CNV deletions include epicanthus, microcephaly, and seizures, while individuals with *YBX3* CNV deletions often exhibit strabismus and scoliosis. However, as is the case with the majority of CNVs, all *YBX-*containing CNV deletions included other genes that may be contributing to these individuals’ phenotypes.

**Figure 4:**
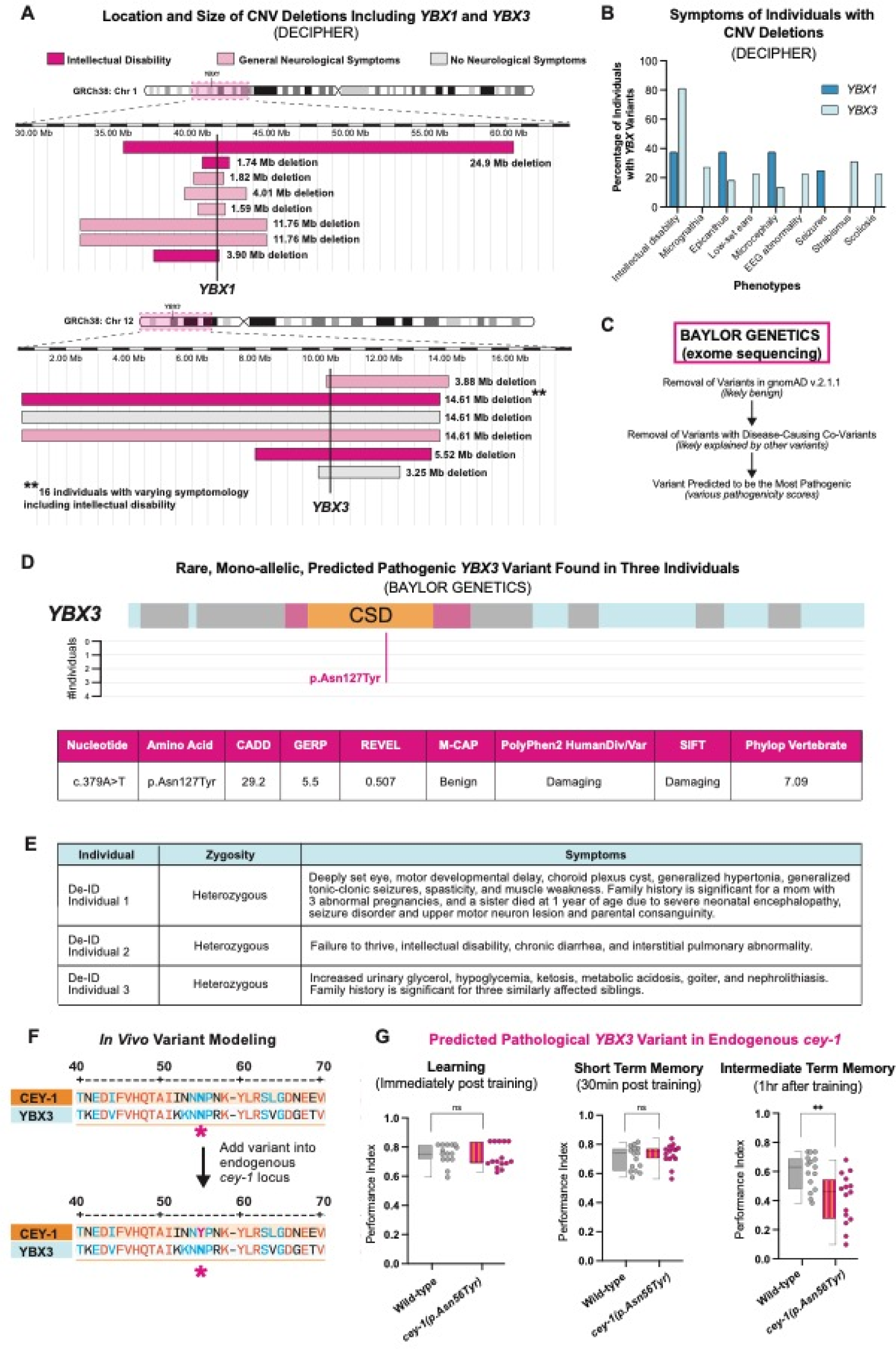
Copy number variants in *YBX1* and *YBX3* found in individuals with neurological symptoms and *in vivo* modeling of a predicted pathogenic single nucleotide variant in YBX3. (**A**) Location, size, and neurological symptoms associated with copy number variant (CNV) deletions reported in DECIPHER. The exact location of *YBX1* and *YBX3* are shown with a bold vertical line. The area around the gene affected by the CNV deletions is highlighted with a pink box on their respective chromosome ideograms at the top of the figure. CNVs where the individual had neurological symptoms are in light pink, and CNVs where the individual had associated intellectual disability are in dark pink. (**B**) Intellectual disability is the most often associated symptom of patients with CNV deletions of *YBX1* or *YBX3* based on DECIPHER. (**C**) Workflow to examine single nucleotide variants (SNVs) that may be disease-linked using the postnatal exome sequencing database from Baylor Genetics. Several strategies were used to remove likely non-pathogenic variants, including removing cases with known co-variants and removal of variants found in the general population (gnomAD v2.1.1)[38]. (**D**) Location and pathogenicity score of rare a heterozygous SNV in *YBX3* found in three individuals, two with indicated neurological symptoms. The variant we identified, p.Asn127Tyr, had the highest predicted pathogenicity using seven different metrics (see text). (**E**) Table of p.Asn127Tyr cases from Baylor Genetics de-identified exome sequencing dataset. Two of the three individuals have neurological symptomology reported, such as intellectual disability and seizures. (**F**) Location of p.Asn127Tyr variant shows conserved residues between CEY-1 and YBX3 and location of variant when introduced into the endogenous CEY-1 locus. (**G**) Introduction of p.Asn127Tyr into the endogenous *cey-1* locus (p.Asn56Tyr) causes ITM deficits. Box and whisker plot as in Figure 1. n = 15 per genotype. **p<0.01, ns, not significant (p>0.05).

In the case of *YBX1*, the minimal overlap between CNV losses affecting the *YBX1* loci is 1.08 Mb containing 34 genes, including two loci that are predicted to be loss of function intolerant: forkhead box J3 (*FOXJ3*)[103] and solute carrier family 2 member 1 (*SLC2A1*) [104]. *SLC2A1* is associated with autosomal dominant Dystonia 9 [105,106], GLUT1 deficiency syndrome 1 [107,108], GLUT1 deficiency syndrome 2 [109,110], Stomatin-deficient cryohydrocytosis with neurologic defects [111,112], and susceptibility to idiopathic generalized epilepsy 12 [113,114]. In the case of *YBX3*, the minimal overlap between CNV deletions affecting the *YBX3* loci is 3.24 Mb containing 68 genes, including three loci that are predicted to be loss of function intolerant: ETS variant transcription factor 6 (*ETV6*)[115], low density lipoprotein receptor-related protein 6 (*LRP6*)[116], and dual-specificity phosphatase 16 (*DUSP16*)[117]. However, unlike the loci surrounding *YBX1*, none of the loss of function intolerant loci around *YBX3* are known to cause monogenetic neurological disease. Summary statistics for CNV DECIPHER data for *YBX1* can be found in **Figure S7** and for *YBX3* in **Figure S8**. All individuals with CNV duplications and deletions in *YBX1* or *YBX3* as well as their DECIPHER phenotypic data can be found in **Table S1.**

Taken together, these data suggest that loss of function CNVs including the *YBX1* and *YBX3* loci may be associated with neurological dysfunction; however, the confounding variable of multi-gene deletions prevents any direct association. As a result, we decided to model a rare single nucleotide variant in *YBX3* for biological function *in vivo*.

### Introducing a predicted deleterious variant in *YBX3* into the endogenous *cey-1* locus causes memory deficits *in vivo*

While publicly available CNV datasets suggest that single-copy loss of *YBXs* may potentially contribute to neurological symptoms, the minimal overlap regions for CNV deletions containing *YBX1* and *YBX3* include loci beyond the gene of interest that are associated with monogenic neurological disorders in humans. Therefore, single-nucleotide variants (SNVs) in the genes of interest may better link deleterious variants in single genes directly to neurological disorders [10,12,118–120] compared to CNVs with multi-gene deletions.

We received a deidentified dataset of *YBX* variants identified through exome and genome sequencing completed at Baylor Genetics [37] to identify potentially deleterious SNVs in the *YBXs*. To identify variants of interest, we curated rare, heterozygous variants in postnatal individuals, as these are the most likely to be deleterious [121,122]. We first removed SNVs that were most likely not contributing to early developmental brain disorders by eliminating variants found in gnomAD v2.1.1 as well as cases where the subject’s symptomology was explained by co-variants in the genome (**Figure 4C**). We then used seven different pathogenicity metrics to determine which variants were the most likely to be deleterious, including CADD [43,44], GERP [42], REVEL [45], M-CAP [39], PolyPhen 2 [41], SIFT [40], and Phylop Vertebrate [46]. The variant with the highest overall pathogenicity scores was in *YBX3*: c.379A>T resulting in the amino acid change p.Asn127Tyr. The affected amino acid is in the Cold-Shock Domain of the YBX3 protein and, significantly, this variant is not reported in gnomAD v.2.1.1 nor gnomAD v4, suggesting it is rare (**Figure 4D**). We found this variant in three different individuals, two of which present neurological symptoms such as intellectual disability and seizures (**Figure 4E**).

The p.Asn127 residue is highly conserved across species, including between *C. elegans* CEY-1 and human YBX3 (**Figure 4F**). Therefore, we decided to test the potential significance of this variant *in vivo* in *C. elegans* by introducing it into the endogenous *cey-1* locus and examining its effect on memory. We used CRISPR to introduce the same mutation into the endogenous *cey-1* gene at the same conserved residue as in YBX3: p.Asn56Tyr (*cey-1 p.Asn56Tyr (p.Asn56YTyr)).* We then tested for memory deficits at learning, STM, and ITM. Remarkably, introduction of the p.Asn56Tyr variant into *cey-1* caused ITM deficits, suggesting the variant may have consequences on human neurological function (**Figure 4G**). Importantly, CEY-1 p.Asn56Tyr worms had no butanone sensing deficits or motility impairments (**Figure S9**), suggesting that in this case, neurological dysfunction is memory-specific. We performed qRT-PCR and verified that while the *cey-1(rrr12)* allele is a true knockout that undergoes nonsense mediated decay[34], our p.Asn56Tyr variant has no significant effect on levels of *cey-1* mRNA, suggesting this SNV affects protein function rather than gene expression levels (**Figure S9**). By using the worm to model one predicted damaging variant in *YBX3*, our findings suggest that *YBX3* may be of interest for human disease associations. Therefore, we decided to take our rare variant modeling a step further by using humanized worm strains expressing *YBX1/3*.

### *YBX1* and *YBX3* functionally replace *cey-1* in a humanized model and introduction of a predicated deleterious variant into *YBX3* causes memory deficits *in vivo*

Recent studies have highlighted the utility of humanized worm lines in phenotypic modeling of rare human gene variants *in vivo* [123–126]. We decided to use this CRISPR-based approach to determine whether the predicted damaging variant YBX3 p.Asn127Tyr, which we found in *YBX3* in two human individuals with neurological symptoms and one individual without neurological symptoms, affects YBX3 function in the context of memory. We created two humanized worm lines expressing either human *YBX1* or *YBX3* at the endogenous *cey-1* locus, thereby replacing *cey-1* (**Figure 5A**). In both cases, the human orthologs were able to functionally replace *cey-1*: learning, STM, and ITM were comparable to that of wild-type animals (**Figure 5B, 5C, Figure S10**). Intriguingly, worms expressing *YBX3* displayed behavior that was nearly indistinguishable from wild-type animals, exhibiting normal forgetting at two hours post-training, while worms expressing *YBX1* still maintained a modest memory two hours post-training, suggesting potential differences in regulation during active forgetting (**Figure S10**). Given *YBX3* functionally replaces *cey-1* in memory, we introduced the rare predicted deleterious variant p.Asn127Tyr into the humanized *YBX3* locus to determine if the variant would affect memory. Indeed, worms expressing *YBX3(p.Asn127Tyr)* have severe intermediate term memory deficits (**Figure 5D**), mirroring our findings with the equivalent variant inserted into the endogenous *cey-1* locus (**Figure 4G**). None of the humanized strains had impaired butanone sensing or motility, suggesting these deficits are memory-specific (**Figure S10**). Taken together, our findings suggest a high degree of functional conservation between *cey-1* and *YBX3*.

**Figure 5:**
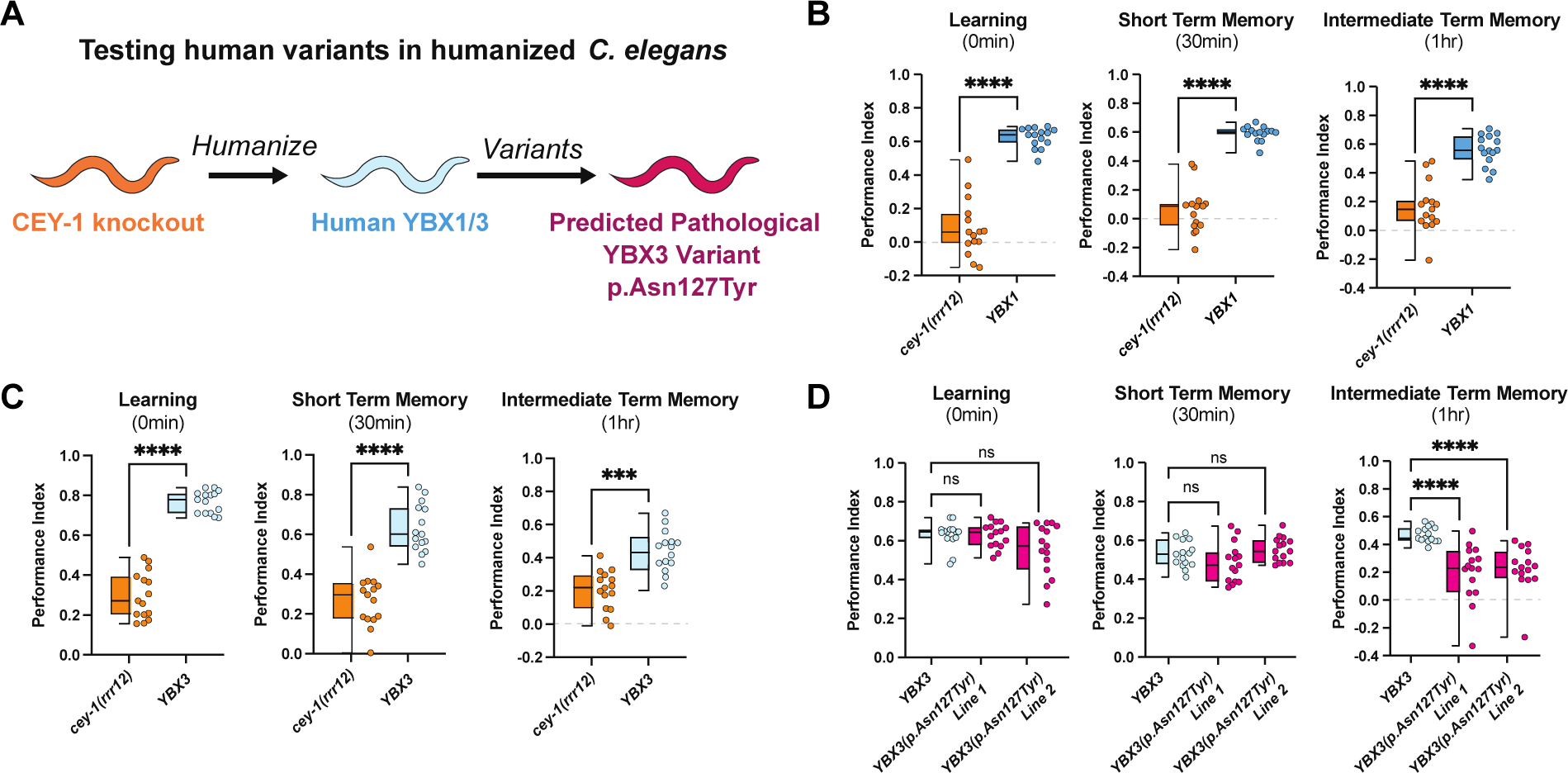
Human *YBX1* and *YBX3* functionally replace *cey-1* in memory *in vivo* and insertion of a rare SNV into *YBX3* disrupts ITM. (**A**) Diagram of workflow for humanizing *C. elegans* to express either human *YBX1* or human *YBX3*, then inserting a rare SNV p.Asn127Tyr into *YBX3*. (**B**) Insertion of human *YBX1* into the endogenous *cey-1* locus rescues learning and memory deficits seen in *cey-1* knockout animals *(cey-1(rrr12))* back to wild-type levels (also see **Figure S10**). (**C**) Insertion of human *YBX3* into the endogenous *cey-1* locus rescues learning and memory deficits seen in *cey-1* knockout animals *(cey-1(rrr12))* back to wild-type levels (also see **Figure S10**). (**D**) Introduction of p.Asn127Tyr into humanized *YBX3* locus causes ITM deficits. Box and whisker plot as in Figure 1. n = 15 per genotype. ****p<0.0001, ns, not significant (p>0.05).

## DISCUSSION

### *C. elegans* as a discovery platform for novel associative memory regulators

We performed an unbiased screen of 20 synaptically-enriched RBPs examining their roles in learning and memory and identified eight novel associative memory regulators, each controlling memory differently. While we focused on the CEY RBP family in this work, five other RBPs remain to be investigated. We found that *rbm-3.1* is required for STM, *rnp-5* is required for STM and ITM, *rpl-28* is required for STM and LTM, *rps-16* is required for STM, ITM, and LTM, and *eef-1B.2* suppresses STM. *rbm-3.1,* ortholog of CIRP, is reported to mediate alcohol-induced spatial memory deficits in mice [127] but has never been associated with memory independently of alcohol use prior to this study. In addition, *eef-1B.2* and *rnp-5* are associated with human neurological disease. Pathogenic variants in EEF1B.2 cause autosomal recessive intellectual disability [128], while *rnp-5* ortholog RNPS1 is a risk factor for both intellectual disability and autism [129,130]. To our knowledge, neither *rps-16* nor *rpl-28* have been studied in the nervous system of any organism.

Here, we demonstrated novel roles for eight RBPs in memory using *C. elegans*. Our results suggest that these RBPs play a conserved role in memory that remains understudied, and future studies should combine worm behavior with human genetic data to examine these and other uncharacterized genes. Taken together, our study highlights the utility of *C. elegans* as a high-throughput screening tool for novel associative memory regulators, including those that control different, molecularly distinct forms of memory.

### Three of the four CEY RBPs control memory

We identified three members of the CEY RBP family, orthologs of the mammalian YBX RBPs, as novel memory molecules. We found that each CEY RBP has its own distinct role in memory: *cey-1* regulates STM/ITM, *cey-2* regulates ITM, and *cey-3* regulates LTM. Of note, *cey- 4* does not appear to be required for memory ability but is expressed throughout the adult nervous system [73,89,90], and therefore likely controls other neuronal functions. The unique role of each CEY in memory is not explained by their expression patterns, as *cey-1* and *cey-4* are both broadly expressed in the nervous system while *cey-2* and *cey-3* are found in very few neurons [73,89,90]. While evidence from the germline suggests the CEY RBPs are promiscuous in binding and are not affiliated with specific mRNA subsets [34], it is unclear whether there is more mRNA target specificity in the nervous system. Future studies examining how each CEY RBP controls the translational landscape of neurons are warranted to determine the independent mechanisms by which the CEYs proteins regulate memory.

### CEY-1 is a novel neuronal memory promoting RBP

Our study of CEY-1, the closest *C. elegans* ortholog to the mammalian YBX proteins, revealed that it acts in the nervous system to promote memory, as neuron-specific rescue of CEY-1 ameliorates behavioral defects cause by the loss of *cey-1,* and an additional copy of CEY-1 only in the nervous system improves memory performance. Future investigations are needed to unveil the molecular mechanisms by which CEY-1 regulates memory, but based on known CEY/YBX biology, we hypothesize that CEY-1 controls transcription and/or translation of mRNAs involved in memory.

YBX1 is reported to bind to plasticity-associated mRNAs in an activity-dependent manner *in vitro* [79], and both CEY-1/YBX proteins regulate polysome formation and translation [34,81,82]. Furthermore, in non-neuronal tissues, YBX1 is reported to interact with known regulators of plasticity-dependent translation, specifically mTOR, AKT and translation initiation machinery [71,81,131]; therefore, it is likely that CEY-1/YBX is involved in these processes as well. Unlike other RBPs linked to plasticity-dependent translation, such as FMRP [132], YBX proteins are relatively non-specific with regards to target mRNAs [82,133]. It is unclear if a subset of molecules required for memory are regulated by YBX, or if it is involved in controlling broad machinery involved in plasticity and memory such as general protein synthesis. Future work will be necessary to determine the exact mechanisms by which CEY-1 promotes memory.

### YBX3 p.Asn127Tyr decreases intermediate term memory in a humanized *C. elegans* model

Given the lack of experimental evidence linking the mammalian YBX proteins to memory, we queried human variant datasets to explore potential associations between YBX dysfunction and neurological symptoms. We focused on *YBX1* and *YBX3* because they are expressed in the adult nervous system, and found that several large CNVs involving either the *YBX1* or *YBX3* loci, as well as one rare missense YBX3 variant, are present in individuals with neurological symptoms including intellectual disability. However, the CNV analysis is limited by other disease-associated genes in the minimal overlap regions involving *YBX1* and *YBX3* and the SNV analysis is limited by the sample size and available de-identified clinical findings.

To assess the biological significance of rare, mono-allelic SNVs in either *YBX1* or *YBX3*, we conducted *in vivo* modeling of the variant with the highest pathogenicity scores. The YBX3 p.Asn127Tyr variant was identified in three individuals, two of whom exhibit neurological symptoms and one individual without neurological symptoms. The p.Asn127Tyr variant is located within the YBX3 CSD at a highly conserved site [85,86,133]. Introduction of the corresponding p.Asn56Tyr variant into the endogenous *cey-1* gene caused ITM deficits. We then decided to perform the same variant study in the context of human *YBX3*. Both *YBX1* and *YBX3* are able to functionally replace *cey-1*, as substitutions using either gene at the *cey-1* locus does not impair learning and memory. Introducing the YBX3 p.Asn127Tyr variant into the humanized *C. elegans* line caused similar ITM deficits as the introduction of the variant into endogenous *cey-1*. Overall, these results suggest three key findings: one, that *cey-1* and *YBX3* are functionally conserved, two, that these data implicate both *YBX1* and *YBX3* in memory for the first time, and three, that this SNV in *YBX3* may be of interest for future studies exploring the role of *YBX3* in human disease.

We believe that the observed ITM deficits observed from the introduction of the p.Asn127Tyr SNV may stem from dysregulation of mRNA translation by CEY-1/YBX3. The CSD, responsible for RNA binding, often lacks a highly specific affinity for a particular RNA motif [86,133]. As a result, proteins with CSDs are often considered master regulators of the mRNA landscape. It is likely that introduction of a variant into the CSD disrupts broad RNA regulation by CEY-1/YBX3, including memory-regulating mRNAs. Regulation of target mRNAs by CEY- 1/YBX3 may be at the level of transcription, translation, or another mechanism, so it is unclear how exactly this variant affects CEY function. Though the mechanism is currently unknown, our data indicate that rare *YBX3* variants may be biologically significant and potentially affect neurologic function.

### Foundations for mammalian *in vivo* studies of YBX proteins in plasticity and neurological disease

Our finding of functional conservation between *C. elegans cey-1* and human *YBX1* and *YBX3* with regards to memory underscores the importance of future mammalian studies to determine the role of the *YBXs* in the nervous system. To date, studying the function of specific YBX proteins in mammalian models is challenging due to the potential for YBXs to genetically compensate for one another and the low survival rate of double knock-outs (Evans *et al.*, 2020). Here, we report that either depleting CEY-1/YBX levels specifically in adulthood or introducing a point variant detected in three human individuals disrupts memory. Each of these manipulations is less likely to be lethal or induce genetic compensation than complete loss during development. Future studies employing similar approaches in mammalian models will be extremely valuable. Moreover, future studies will need to be considerate of loss of function mechanisms of variants. *YBX1* is predicted to be loss of function intolerant (pLI = 1, gnomAD v4.0), suggesting haploinsufficient variants may be deleterious. Conversely, *YBX3* is predicted to be loss of function tolerant (pLI = 0, gnomAD v4.0), suggesting haplosufficiency. Therefore, for rare variants in *YBX3* to cause dysfunction, these variants may be dominant-negative or result in gain-of-function. Future identification of rare variants in the *YBX*s using comprehensive exome or genomic human datasets and modeling variants *in vivo* in mammals will enhance our understanding of mechanisms by which YBX dysfunction may play a role in human disease.

Taken together, our results uncover the CEY-1/YBX RBPs as novel associative memory regulators and potentially new contributors to rare neurological disease. Our current findings are limited to the functional testing of a single variant in *YBX3*. However, further examination of this protein class will provide new insights into the molecular mechanisms of memory and deepen our understanding of how disruption of RBPs contributes to disorders of the nervous system.

## Author Contributions

Conceptualization, ANH and RNA; Methodology, ANH and RNA; Formal Analysis, ANH, RNA, JAR, and HTC; Investigation, ANH, KLB, PRM, PV, EJL, EWP, ESG, and CWN; Resources, ANH and RNA; Writing – Original Draft, ANH and RNA; Writing – Review & Editing, ANH, KLB, ESG, EWP, CWN, JAR, HTC, RNA; Visualization, ANH and RNA; Supervision, ANH, JAR, HTC, RNA; Project Administration, ANH and RNA; Funding Acquisition, ANH and RNA.

## Acknowledgements

We thank the Ciosk lab for generously sharing the cey-1 #1087 strain; the CGC for strains; Ben Jussila and In Vivo Biosystems for strains and helpful discussion; Peter Boag for helpful discussion; and members of the Arey lab, particularly Katie L Brandel and Catherine Stuart for their detailed feedback on the manuscript. We thank Drs. Jill Rosenfeld and Hongzheng Dai from Baylor Genetics for confirming the presence and relevance of the YBX3 p.Asn127Tyr variant.

This study makes use of data generated by the DECIPHER community. A full list of centres who contributed to the generation of the data is available from https://deciphergenomics.org/about/stats and via email from. DECIPHER is hosted by EMBL-EBI and funding for the DECIPHER project was provided by the Wellcome Trust (WT223718/Z/21/Z).

