## Supplemental Figures and Legends for "Behavioral screening of conserved RNA-binding proteins reveals CEY-1/YBX RNA-binding protein dysfunction leads to impairments in memory and cognition"

**Figure S1**

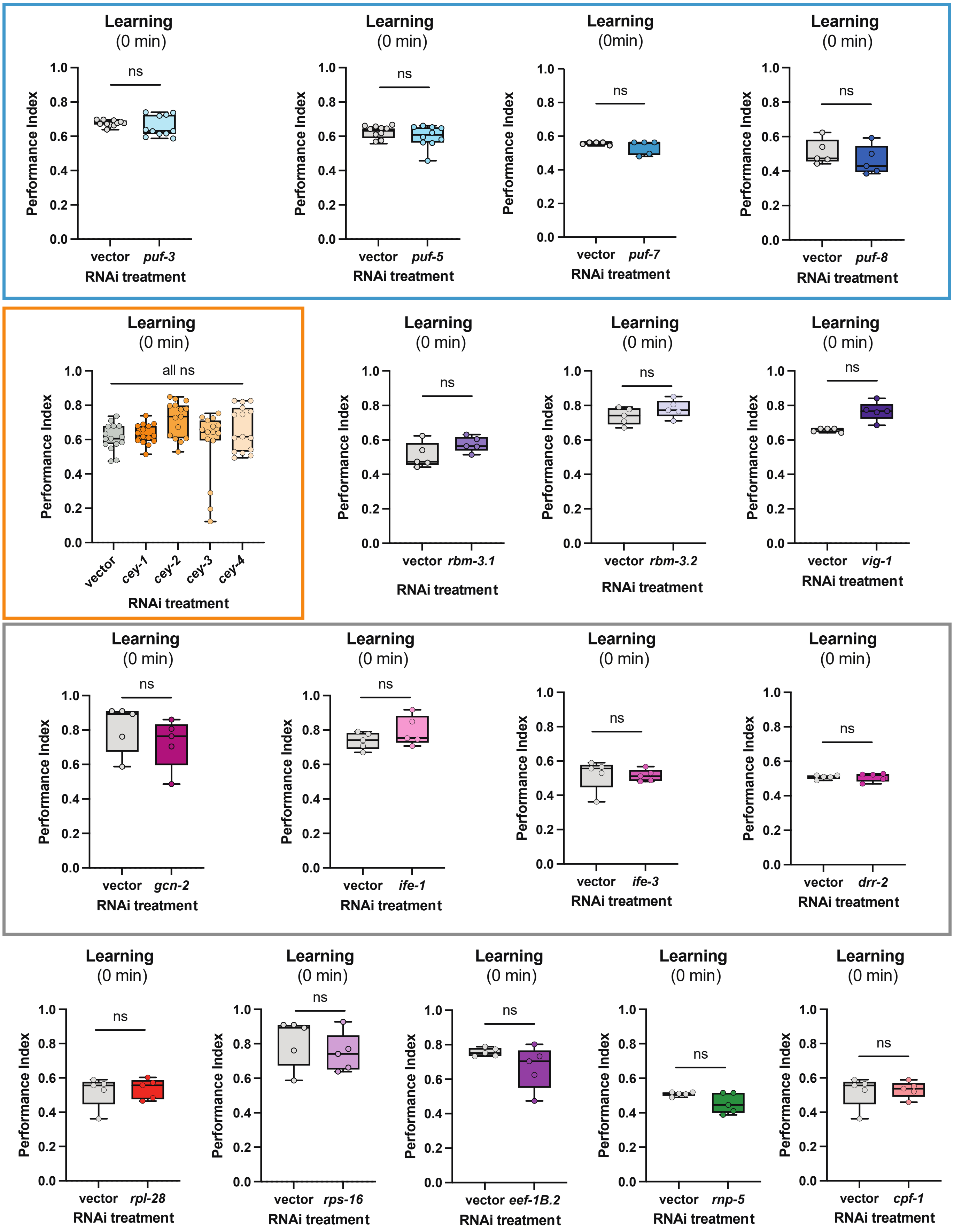
Learning
(0 min)
ns
Performance Index
1.0
0.8
0.6
0.4
0.2
0.0
vector
puf-3
RNAi treatment
Learning
(0 min)
ns
vector
puf-5
RNAi treatment
Learning
(0min)
ns
vector
puf-7
RNAi treatment
Learning
(0 min)
ns
vector
puf-8
RNAi treatment
Learning
(0 min)
all ns
vector
cey-1
cey-2
cey-3
cey-4
RNAi treatment
Learning
(0 min)
ns
vector
rbm-3.1
RNAi treatment
Learning
(0 min)
ns
vector
rbm-3.2
RNAi treatment
Learning
(0 min)
ns
vector
vig-1
RNAi treatment
Learning
(0 min)
ns
vector
gcn-2
RNAi treatment
Learning
(0 min)
ns
vector
ife-1
RNAi treatment
Learning
(0 min)
ns
vector
ife-3
RNAi treatment
Learning
(0 min)
ns
vector
drr-2
RNAi treatment
Learning
(0 min)
ns
vector
rpl-28
RNAi treatment
Learning
(0 min)
ns
vector
rps-16
RNAi treatment
Learning
(0 min)
ns
vector
eef-1B.2
RNAi treatment
Learning
(0 min)
ns
vector
rnp-5
RNAi treatment
Learning
(0 min)
ns
vector
cpf-1
RNAi treatment

Figure S2

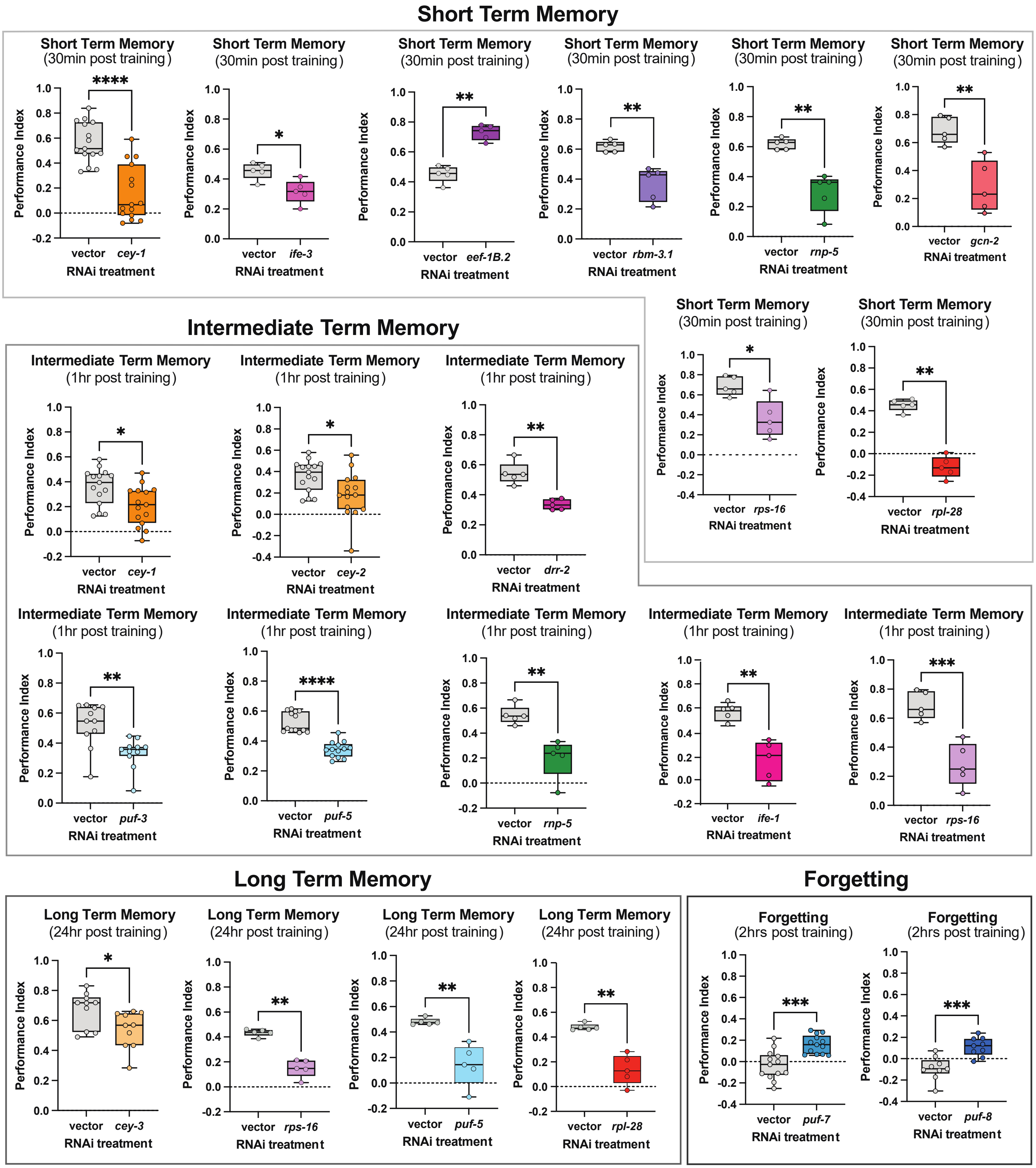

Short Term Memory
Short Term Memory (30min post training)
Performance Index
****
vector cey-1
RNAi treatment
Short Term Memory (30min post training)
*
vector ife-3
RNAi treatment
Short Term Memory (30min post training)
**
vector eef-1B.2
RNAi treatment
Short Term Memory (30min post training)
**
vector rbm-3.1
RNAi treatment
Short Term Memory (30min post training)
**
vector rnp-5
RNAi treatment
Short Term Memory (30min post training)
**
vector gcn-2
RNAi treatment
Short Term Memory (30min post training)
*
vector rps-16
RNAi treatment
Short Term Memory (30min post training)
**
vector rpl-28
RNAi treatment
Intermediate Term Memory
Intermediate Term Memory (1hr post training)
*
vector cey-1
RNAi treatment
Intermediate Term Memory (1hr post training)
*
vector cey-2
RNAi treatment
Intermediate Term Memory (1hr post training)
**
vector drr-2
RNAi treatment
Intermediate Term Memory (1hr post training)
**
vector puf-3
RNAi treatment
Intermediate Term Memory (1hr post training)
****
vector puf-5
RNAi treatment
Intermediate Term Memory (1hr post training)
**
vector rnp-5
RNAi treatment
Intermediate Term Memory (1hr post training)
**
vector ife-1
RNAi treatment
Intermediate Term Memory (1hr post training)
***
vector rps-16
RNAi treatment
Long Term Memory
Long Term Memory (24hr post training)
*
vector cey-3
RNAi treatment
Long Term Memory (24hr post training)
**
vector rps-16
RNAi treatment
Long Term Memory (24hr post training)
**
vector puf-5
RNAi treatment
Long Term Memory (24hr post training)
**
vector rpl-28
RNAi treatment
Forgetting
Forgetting (2hrs post training)
***
vector puf-7
RNAi treatment
Forgetting (2hrs post training)
***
vector puf-8
RNAi treatment

### Figure S3

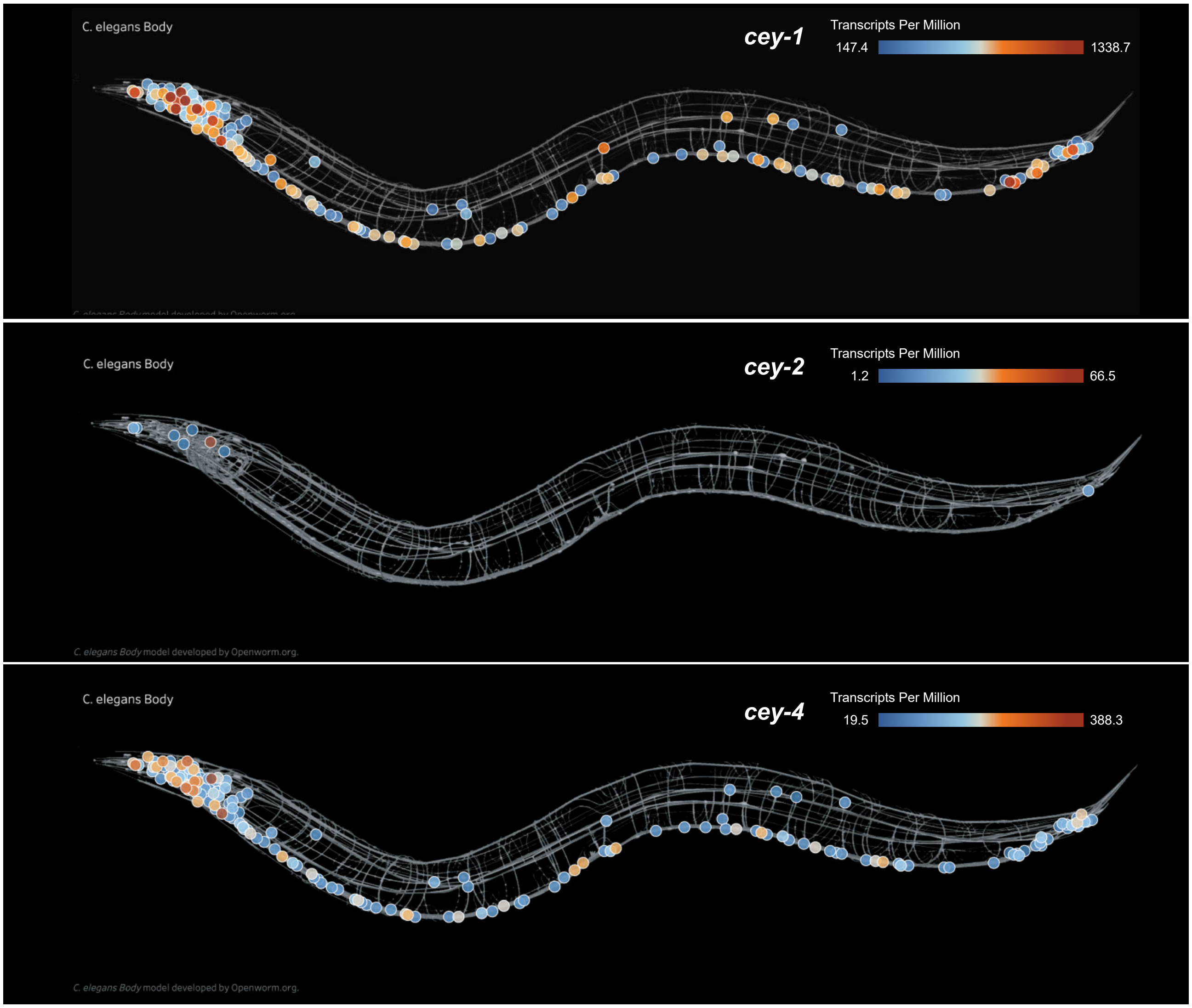
C. elegans Body
cey-1
Transcripts Per Million
147.4
1338.7
C. elegans Body
cey-2
Transcripts Per Million
1.2
66.5
C. elegans Body model developed by Openworm.org.
C. elegans Body
cey-4
Transcripts Per Million
19.5
388.3
C. elegans Body model developed by Openworm.org.

**Figure S4**

A

| Publication Dataset | Type of Sequencing | *cey-1* mRNA present? |
|---|---|---|
| *Kaletsky et al. 2018* | Neuron-specific RNA-seq in Day 1 adults | Yes |
| *Yao et al. 2018* | Integrated neuron-specific transcriptomics from over 2400 publications | Yes |
| *Arey et al. 2019* | Nervous system-wide presynaptic RNAseq in Day 1 adults | Yes |
| *Taylor et al. 2021** | Single-cell RNA-seq in neurons of L4 larvae | Yes (42.4%) |
| *Wang et al. 2022* | Neuron-specific Smart-seq2 in Day 1 and Day 8 adults | Yes |
| *St. Ange et al. 2024* | Single nuclei RNAseq of Day 1 adult nervous system | Yes |

**Percentage of neurons transcript is detected in assuming 302 neurons with a stringency of 2*

B

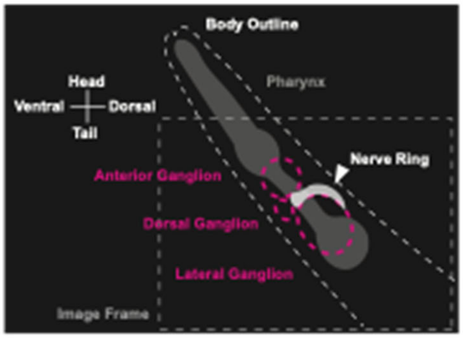


C

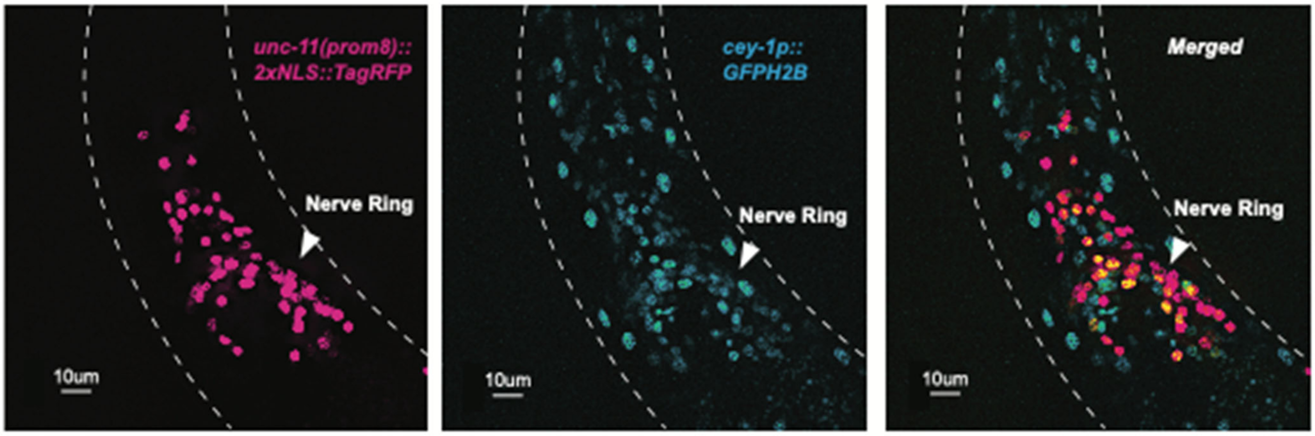

**Figure S5**

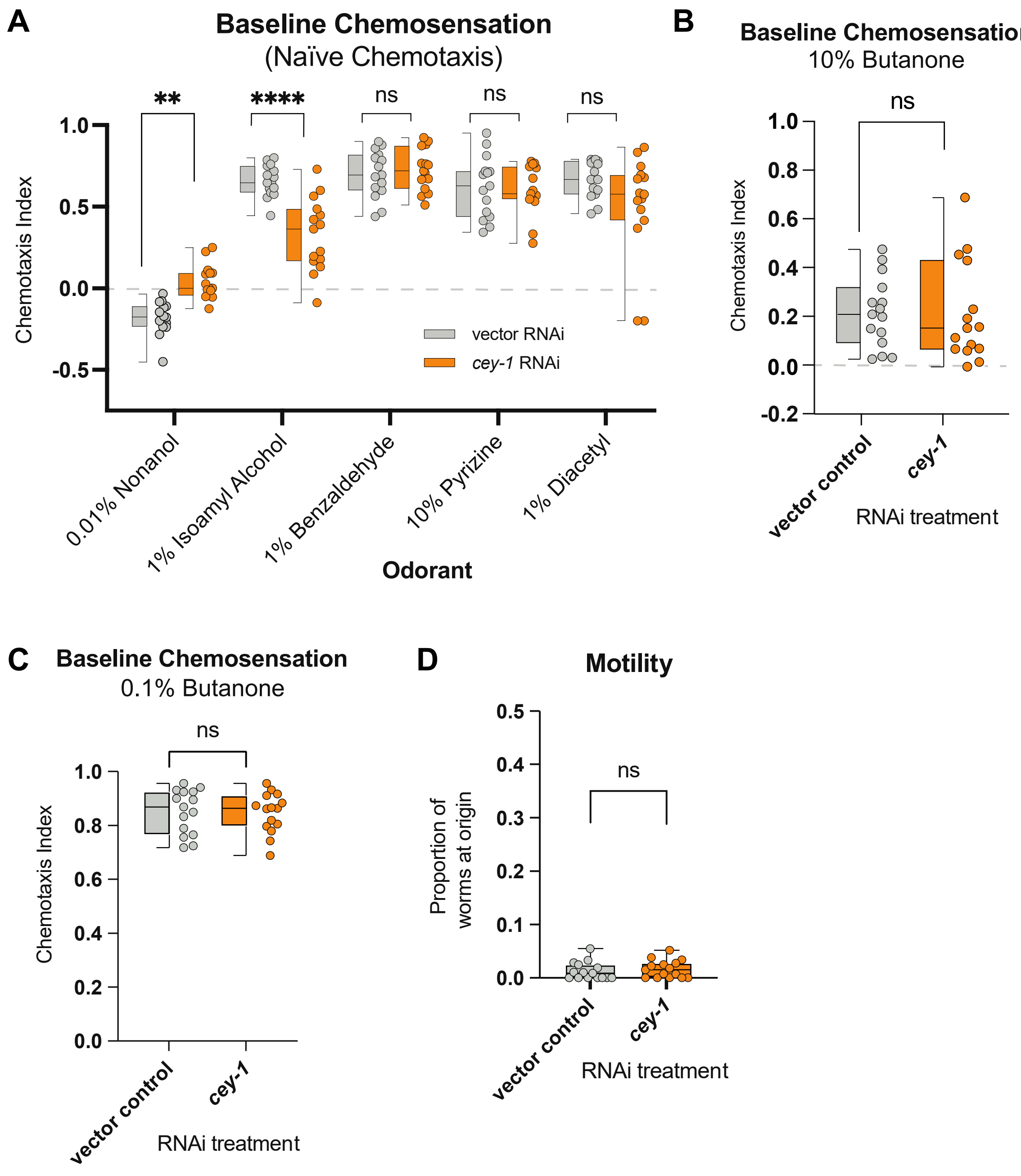
A
Baseline Chemosensation
(Naïve Chemotaxis)
**
****
ns
ns
ns
1.0
0.5
0.0
-0.5
Chemotaxis Index
vector RNAi
cey-1 RNAi
0.01% Nonanol
1% Isoamyl Alcohol
1% Benzaldehyde
10% Pyrizine
1% Diacetyl
Odorant
B
Baseline Chemosensation
10% Butanone
ns
1.0
0.8
0.6
0.4
0.2
0.0
-0.2
Chemotaxis Index
vector control
cey-1
RNAi treatment
C
Baseline Chemosensation
0.1% Butanone
ns
1.0
0.8
0.6
0.4
0.2
0.0
Chemotaxis Index
vector control
cey-1
RNAi treatment
D
Motility
0.5
0.4
0.3
0.2
0.1
0.0
ns
Proportion of worms at origin
vector control
cey-1
RNAi treatment

**Figure S6**

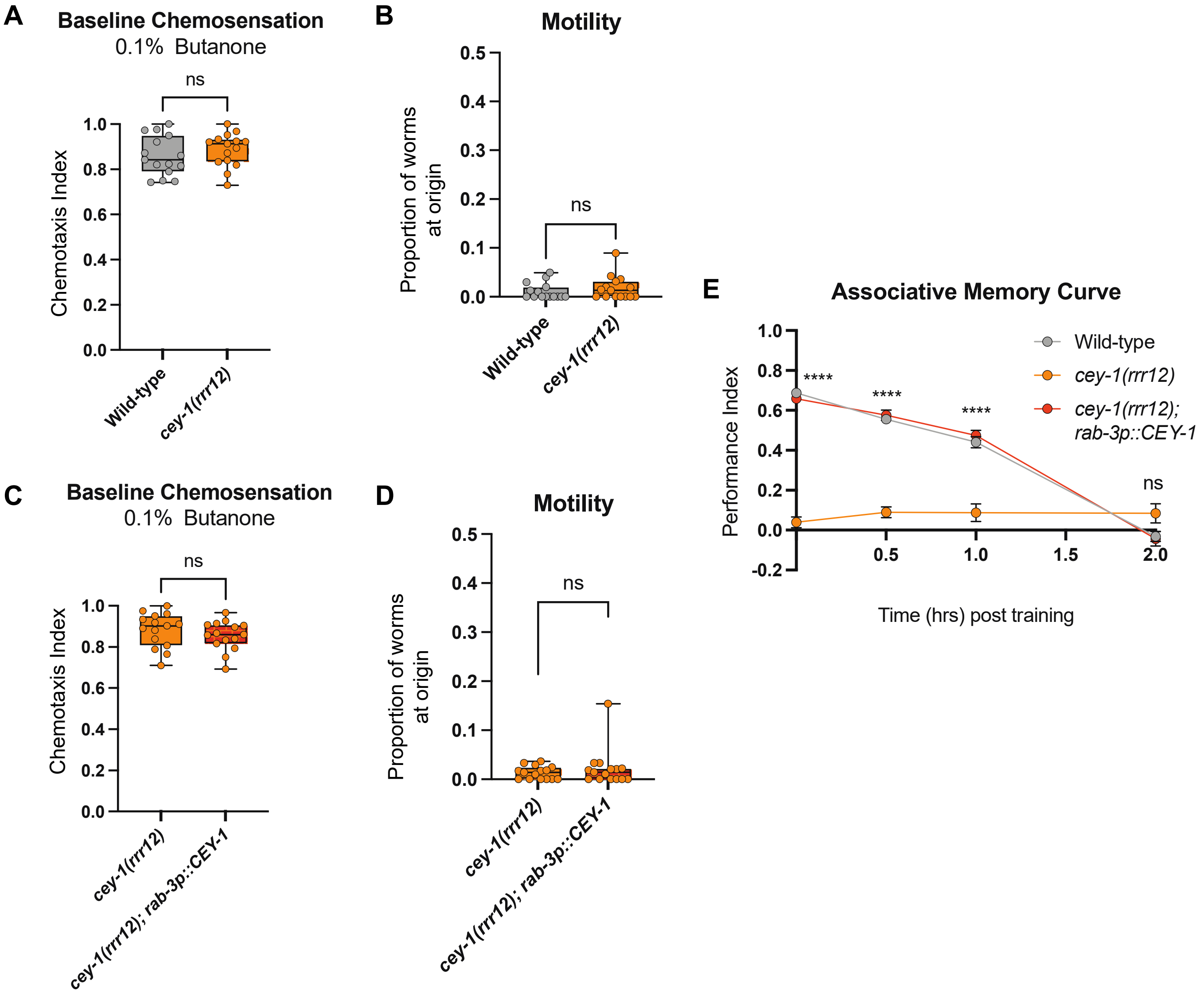
A
Baseline Chemosensation
0.1% Butanone
ns
Chemotaxis Index
1.0
0.8
0.6
0.4
0.2
0.0
Wild-type
cey-1(rrr12)
B
Motility
Proportion of worms at origin
0.5
0.4
0.3
0.2
0.1
0.0
ns
Wild-type
cey-1(rrr12)
C
Baseline Chemosensation
0.1% Butanone
ns
Chemotaxis Index
cey-1(rrr12)
cey-1(rrr12); rab-3p::CEY-1
D
Motility
Proportion of worms at origin
ns
cey-1(rrr12)
cey-1(rrr12); rab-3p::CEY-1
E
Associative Memory Curve
Performance Index
1.0
0.8
0.6
0.4
0.2
0.0
-0.2
****
****
****
ns
0.5
1.0
1.5
2.0
Time (hrs) post training
Wild-type
cey-1(rrr12)
cey-1(rrr12); rab-3p::CEY-1

**Figure S7**

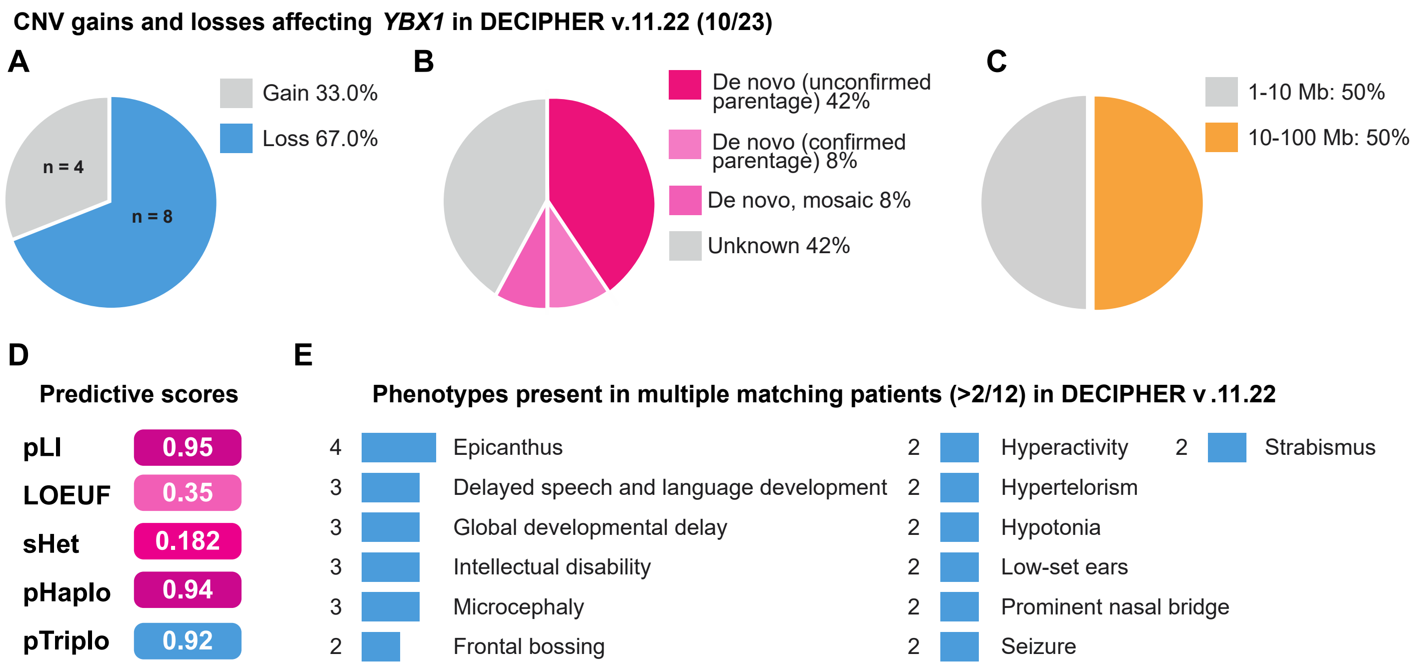
CNV gains and losses affecting *YBX1* in DECIPHER v.11.22 (10/23)
A
Gain 33.0%
Loss 67.0%
n = 4
n = 8
B
De novo (unconfirmed parentage) 42%
De novo (confirmed parentage) 8%
De novo, mosaic 8%
Unknown 42%
C
1-10 Mb: 50%
10-100 Mb: 50%
D
Predictive scores
pLI 0.95
LOEUF 0.35
sHet 0.182
pHaplo 0.94
pTriplo 0.92
E
Phenotypes present in multiple matching patients (>2/12) in DECIPHER v .11.22
4 Epicanthus
3 Delayed speech and language development
3 Global developmental delay
3 Intellectual disability
3 Microcephaly
2 Frontal bossing
2 Hyperactivity
2 Hypertelorism
2 Hypotonia
2 Low-set ears
2 Prominent nasal bridge
2 Seizure
2 Strabismus

### Figure S8

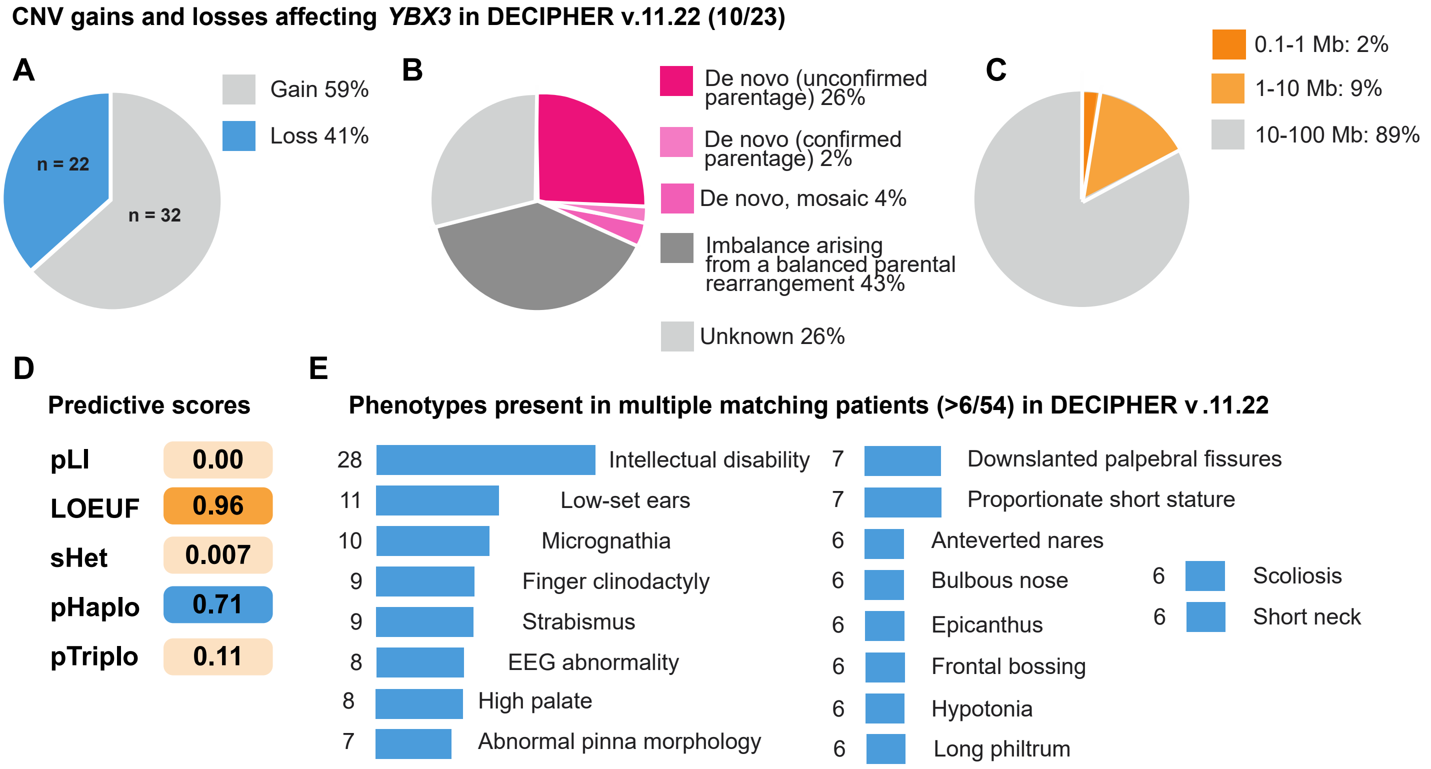

CNV gains and losses affecting *YBX3* in DECIPHER v.11.22 (10/23)
A
Gain 59%
Loss 41%
n = 22
n = 32
B
De novo (unconfirmed parentage) 26%
De novo (confirmed parentage) 2%
De novo, mosaic 4%
Imbalance arising from a balanced parental rearrangement 43%
Unknown 26%
C
0.1-1 Mb: 2%
1-10 Mb: 9%
10-100 Mb: 89%
D
Predictive scores
pLI 0.00
LOEUF 0.96
sHet 0.007
pHaplo 0.71
pTriplo 0.11
E
Phenotypes present in multiple matching patients (>6/54) in DECIPHER v .11.22
28 Intellectual disability
11 Low-set ears
10 Micrognathia
9 Finger clinodactyly
9 Strabismus
8 EEG abnormality
8 High palate
7 Abnormal pinna morphology
7 Downslanted palpebral fissures
7 Proportionate short stature
6 Anteverted nares
6 Bulbous nose
6 Epicanthus
6 Frontal bossing
6 Hypotonia
6 Long philtrum
6 Scoliosis
6 Short neck

**Figure S9**

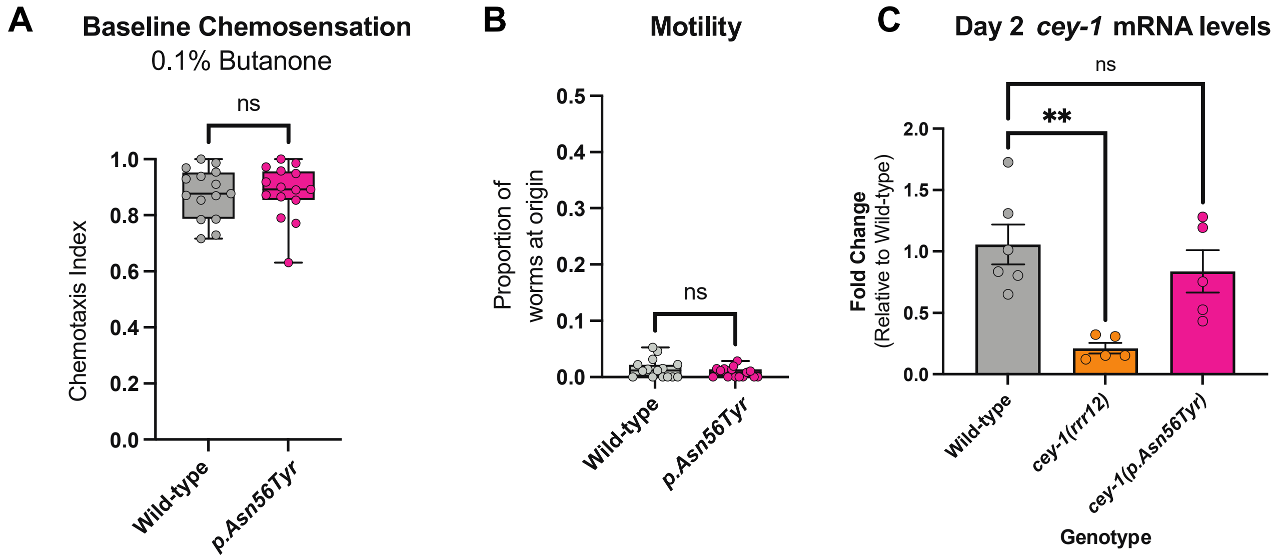

A
Baseline Chemosensation
0.1% Butanone
ns
Chemotaxis Index
1.0
0.8
0.6
0.4
0.2
0.0
Wild-type
p.Asn56Tyr
B
Motility
Proportion of worms at origin
0.5
0.4
0.3
0.2
0.1
0.0
ns
Wild-type
p.Asn56Tyr
C
Day 2 cey-1 mRNA levels
ns
**
Fold Change
(Relative to Wild-type)
2.0
1.5
1.0
0.5
0.0
Wild-type
cey-1(rrr12)
cey-1(p.Asn56Tyr)
Genotype

**Figure S10**

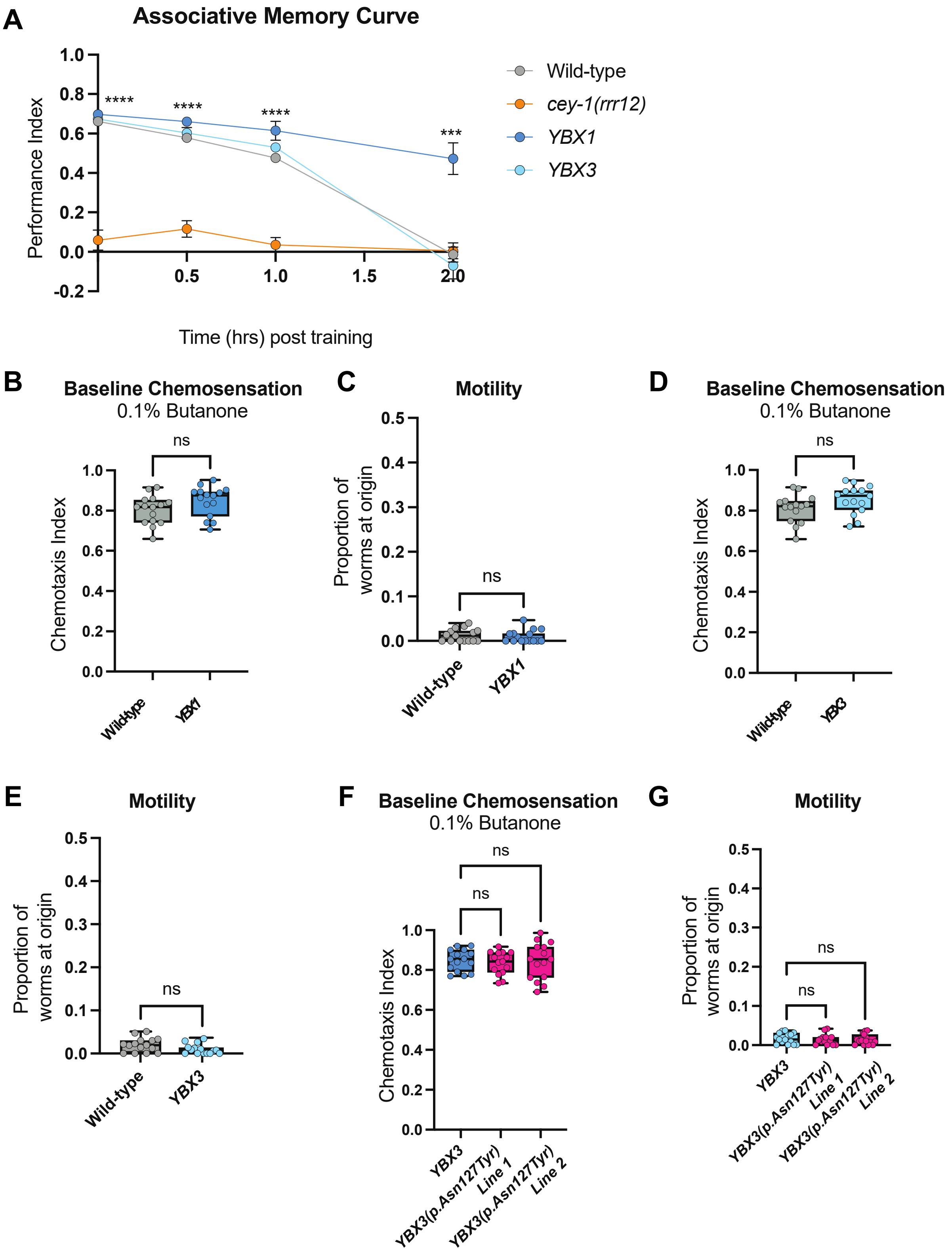

A
Associative Memory Curve
Performance Index
Time (hrs) post training
Wild-type
cey-1(rrr12)
YBX1
YBX3
****
****
****
***
B
Baseline Chemosensation
0.1% Butanone
Chemotaxis Index
ns
Wild-type
YBX1
C
Motility
Proportion of worms at origin
ns
Wild-type
YBX1
D
Baseline Chemosensation
0.1% Butanone
Chemotaxis Index
ns
Wild-type
YBX3
E
Motility
Proportion of worms at origin
ns
Wild-type
YBX3
F
Baseline Chemosensation
0.1% Butanone
Chemotaxis Index
ns
ns
YBX3
YBX3(p.Asn127Tyr) Line 1
YBX3(p.Asn127Tyr) Line 2
G
Motility
Proportion of worms at origin
ns
ns
YBX3
YBX3(p.Asn127Tyr) Line 1
YBX3(p.Asn127Tyr) Line 2

**Supplementary Figure Legends:**

**Figure S1.** RNAi-based knockdown of RBPs does not have any detectable effect on learning ability. Boxes signify 3+ RBPs in the same protein family or class (blue box include PUF RBPs, orange box includes CEY RBPs, grey box includes translation initiation machinery). Box and whisker plots are shown for the learning timepoints from the STM/ITM assays for each RBP screened. Box and whisker plot: the center line denotes the median value (50th percentile) while the box contains the 25th to 75th percentiles. Whiskers mark the 5th and 95th percentiles. ns, not significant ($p>0.05$).

**Figure S2.** Results of unbiased screen of 20 RBPs reveals both known and novel associative memory regulators. All RBPs with significantly altered memory are shown divided by memory timepoint. Box and whisker plot: the center line denotes the median value (50th percentile) while the box contains the 25th to 75th percentiles. Whiskers mark the 5th and 95th percentiles. $n \geq$ 5-10 per RNAi treatment. *$p<0.05$,**$p<0.01$, ***$p<0.001$, ****$p<0.0001$.

**Figure S3.** Images generated from publicly available VISTA (Visualizing the Spatial Transcriptome of the *C. elegans* Nervous System) [1] reveal differences in expression across each of the *cey* RBPs at L4. While *cey-1* and *cey-4* have broad expression, *cey-2* is only located in eight neurons at L4.

**Figure S4.** Data from multiple transcriptomic datasets as well as microscopy images reveal broad expression of CEY-1 in the nervous system of adult worms. (A) Neuron-specific and/or single-cell RNA-seq data compiled from five different publications suggest *cey-1* mRNA is expressed in the adult nervous system [2–8]. (B) Diagram of microscopy images shown in (C). *C. elegans* head is labeled including the location of the pharynx and main neuronal ganglia/nerve rings. (C) A transcriptional *cey-1* reporter suggests the gene is broadly expressed in the neurons in the head at baseline conditions. Representative image of Day 2 adult worms with

RFP-labeled neuronal nuclei (*unc-11(prom8)::2xNLS::TagRFP*)) pseudocolored magenta and a nuclear GFPH2B *cey-1* promoter fusion *(cey1p::GFPH2B*) pseudocolored cyan show colocalization.

**Figure S5.** Knockdown of *cey-1* specifically in the adult nervous system does not impair butanone sensing or motility. (A) Naïve battery of both negative and positive odors reveals deficits in both nonanol and isoamyl alcohol chemotaxis, but not in benzaldehyde, pyrizine, or diacetyl chemotaxis at attractive concentrations. (B) Baseline chemosensation for butanone at neutral concentration of 10% is unaffected by neuron-specific knockdown of *cey-1*. n = 15 per RNAi treatment. (C) Baseline chemosensation for butanone at an attractive concentration of 0.1% is unaffected by neuron-specific knockdown of *cey-1*. n = 15 per RNAi treatment. (D) Motility, measured as proportion of worms at the origin of a chemotaxis plate, is unaffected by neuron-specific loss of *cey-1*. n = 15 per RNAi treatment. Box and whisker plot: the center line denotes the median value (50th percentile) while the box contains the 25th to 75th percentiles. Whiskers mark the 5th and 95th percentiles. **p<0.01, ****p<0.0001. ns, not significant (p>0.05).

**Figure S6.** *cey-1* loss-of-function mutants (*cey-1*(*rrr12*)) as well as worms with neuron-specific rescue of *cey-1* ((*cey-1(rrr12)*; *rab-3p::CEY-1*)) have no deficits in butanone sensing or motility, and neuron-specific rescue of *cey-1* restores learning and memory ability to wild-type levels (A) Baseline chemosensation for butanone at an attractive concentration of 0.001% is unaffected by whole-body loss of *cey-1*. n = 15 per genotype. (B) Motility, measured as proportion of worms at the origin of a chemotaxis plate, is unaffected by whole-body loss of *cey-1*. n = 15 per genotype. (C) Baseline chemosensation for butanone at an attractive concentration of 0.1% is unaffected by neuron-specific rescue of *cey-1*. n = 15 per genotype. (D) Motility, measured as proportion of worms at the origin of a chemotaxis plate, is unaffected by neuron-specific rescue of *cey-1*. n = 15 per genotype. (E) Associative memory curve comparing wild-type, *cey-1(rrr12)* knockout worms, and nervous system specific rescue worms *(cey-1(rrr12);rab-3p::CEY-1)* shows that while knockouts have no associative learning or memory, neuron-specific rescue of *cey-1* allows for learning and memory equivalent to wild-type levels. Box and whisker plot: the center line denotes the median value (50th percentile) while the box contains the 25th to 75th percentiles. Whiskers mark the 5th and 95th percentiles. ns, not significant ($p>0.05$).

**Figure S7.** CNV gains and losses data for *YBX1* in DECIPHER[9]. (A) Percentages of individuals reported with gain or loss CNVs that include *YBX1*. (B) Percentages of mechanisms of inheritance of gain and loss CNVs that include *YBX1*. (C) Size of CNV gain and losses that include *YBX1*. (D) Predictive scores for *YBX1* from gnomAD v.2.11[10] suggest that *YBX1* is intolerant to loss of function variants and is haploinsufficient. (E) Phenotypes of patients specifically with deletion/loss CNVs including *YBX1* include epicanthus, delayed speech and development, intellectual disability, and other neurological features. (F) Individuals reported in DECIPHER with CNV losses including *YBX1*, including the sex of the individual, the size of the deletion, the mechanism of inheritance (if known), the coordinates of the CNV loss, and the individual's reported phenotype.

**Figure S8.** CNV gains and losses data for *YBX3* in DECIPHER[9]. (A) Percentages of individuals reported with gain or loss CNVs that include *YBX3*. (B) Percentages of mechanisms of inheritance of gain and loss CNVs that include *YBX3*. (C) Size of CNV gain and losses that include *YBX3*. (D) Predictive scores for *YBX3* from gnomAD v.2.11[10] suggest that *YBX3* is tolerant to loss of function variants and is haplosufficient, suggesting variants may instead be deleterious by being dominant negative or gain-of-function. (E) Phenotypes of patients specifically with deletion/loss CNVs including *YBX3*, primarily intellectual disability, low-set ears, micrognathia, and other neurological features. (F) Individuals reported in DECIPHER with CNV losses including *YBX3*, including the sex of the individual, the size of the deletion, the mechanism of inheritance (if known), the coordinates of the CNV loss, and the individual's reported phenotype.

**Figure S9.** Introduction of a point mutation from human individuals into *cey-1 (cey-1 (p.Asn56Tyr))* does not cause butanone sensing or motility defects. (A) Baseline chemosensation for butanone at an attractive concentration of 0. 1% is unaffected by the p.Asn56Tyr variant in *cey-1*. n = 15 per genotype. (B) Motility, measured as proportion of worms at the origin of a chemotaxis plate, is unaffected by p.Asn56Tyr variant in *cey-1*. n = 15 per genotype. Box and whisker plot: the center line denotes the median value (50th percentile) while the box contains the 25th to 75th percentiles. Whiskers mark the 5th and 95th percentiles. ns, not significant (p>0.05). (C) qRT-PCR of *cey-1* mRNA levels in Day 2 adults shows that while *cey-1* knockouts *(cey-1(rrr12))* worms undergo nonsense mediated decay[11] resulting in reduced *cey-1* mRNA levels, introduction of the p.Asn56Tyr variant does has no significant effect on *cey-1* expression. n = 6 per genotype. **p<0.01.

**Figure S10.** Humanized lines expressing *YBX1* or *YBX3*, including those with a SNV in *YBX3*, have normal butanone sensing and motility. (A) Associative memory curve comparing wild-type, *cey-1* knockouts, and humanized lines expressing either *YBX1* or *YBX3* at the endogenous *cey-1* locus. While worms expressing *YBX3* have normal memory performance, those expressing *YBX1* appear to have extended memory, as they have a strong association for butanone even two hours post-training. (B) Baseline chemosensation for butanone at an attractive concentration of 0.1% is normal in worms expressing *YBX1*. n = 15 per genotype. (C) Motility, measured as proportion of worms at the origin of a chemotaxis plate, is unaffected by *YBX1*. n = 15 per genotype. (D) Baseline chemosensation is normal in worms expressing *YBX3* at the *cey-1* locus. n = 15 per genotype. (E) Motility is unaffected by *YBX3*. n = 15 per genotype. (F) Baseline chemosensation is unaffected by expressing *YBX3(p.Asn127Tyr)* at the *cey-1* locus. n = 15 per genotype. (G) Motility is unaffected by *YBX3(p.Asn127Tyr)*. n = 15 per genotype.
